# Subsecond whole-brain neural dynamics identified by hidden Markov modeling reflect value-based decision making in humans

**DOI:** 10.64898/2026.05.14.724334

**Authors:** Ryuta Aoki, Kazuki Iijima, Hiroshi Yamada, Kenji Matsumoto, Mitsunari Abe, Takashi Hanakawa, Madoka Matsumoto

## Abstract

Value-based decision making emerges from coordinated neural dynamics across distributed brain networks. Recent studies using noninvasive whole-brain measurements in humans have highlighted the importance of neural activity in the 2–10 Hz frequency band for value-based decision making. Using magnetoencephalography and hidden Markov model (HMM) analysis, we examined whether and how whole-brain neural dynamics in this frequency band, evolving on a timescale of a few hundred milliseconds, reflect value-based decision processes. Thirty-five healthy adults (females and males) made binary choices between risky and sure options. Trial-wise subjective values were estimated using behavioral economic modeling based on prospect theory. We found that HMM-derived trial-by-trial whole-brain neural dynamics (defined by 2–10 Hz amplitude envelopes in distributed brain regions and their interregional coupling) were associated with the subjective values of choice options in a manner distinct from simple perceptual- or motor-evoked activity. Notably, these trial-by-trial whole-brain dynamics covaried with the difference in subjective values between the chosen and unchosen options when the neural data were time-locked to participants’ responses, but not when time-locked to option onset. These findings revealed a crucial link between subsecond whole-brain neural dynamics and trial-by-trial decision variables, providing insights into how value-based decision processes unfold over time in the human brain.

## Introduction

Value-based decision making is a dynamic process involving simultaneous, interrelated computations that unfold over time (Hunt et al., 2015, Hunt and Hayden, 2017). During these computations, various neural signals emerge, including those reflecting the subjective values of choice options (option valuation) and differences in value between options (value comparison). These neural and computational processes give rise to the eventual choice, manifested as the selection of a specific action (Rangel et al., 2008, Kable and Glimcher, 2009, Basten et al., 2010, Williams et al., 2021). Choices are often followed by anticipatory and evaluative processes related to choice outcomes before and after the delivery of outcome feedback (Delgado et al., 2000, McCoy and Platt, 2005, Matsumoto et al., 2007, Rangel et al., 2008, Matsumoto et al., 2022). A large body of evidence from human neuroimaging and animal electrophysiological studies has shown that these decision-making processes are underpinned by diverse brain regions (Rushworth et al., 2009, Rangel and Hare, 2010, Hunt and Hayden, 2017). Existing research has emphasized the central role of the fronto-striatal network (striatum and medial prefrontal cortex [mPFC]) in the valuation of choice and feedback (Delgado, 2007, Levy and Glimcher, 2012, Bartra et al., 2013, Clithero and Rangel, 2014), linking this network to the dopaminergic system and the brain’s valuation system (O’Doherty and Bossaerts, 2008, Lebreton et al., 2009, Haber and Knutson, 2010). However, prior work has also demonstrated the key roles of other brain regions, including the lateral PFC (Matsumoto et al., 2003, Hare et al., 2009, Rushworth et al., 2011, Hunt et al., 2018, Hunt, 2021), anterior insula (Weller et al., 2009, Clark et al., 2014, Knutson and Huettel, 2015), anterior cingulate cortex (Holroyd and Yeung, 2012, Shenhav et al., 2013, Alexander and Brown, 2019), posterior cingulate cortex/precuneus (Bartra et al., 2013, Krug et al., 2014), parietal cortex (Platt and Glimcher, 1999, Huettel et al., 2005, Platt and Huettel, 2008), and hippocampus (Wimmer and Shohamy, 2012, Bakkour et al., 2019, Liu et al., 2019). These findings indicate that value-based decision making emerges from the orchestration of neural activity across the entire brain. Recent functional magnetic resonance imaging (fMRI) studies have further supported this view, showing that subjective value is represented by multivoxel patterns distributed across the whole brain, not just by activation in a few regions traditionally associated with value (Vickery et al., 2011, Chang et al., 2022, Kragel et al., 2023). Although these fMRI studies have advanced our understanding of the brain-wide neural signatures of subjective value, the temporal resolution of fMRI (typically ∼1 s) is insufficient to capture whole-brain dynamics that evolve in the order of hundreds of milliseconds (Baker et al., 2014, Quinn et al., 2018). Thus, the temporal profiles of subsecond whole-brain dynamics associated with value-based decision making remain unclear.

Magnetoencephalography (MEG) enables noninvasive recording of electromagnetic brain activity, and is particularly suitable for studying rapid (i.e., subsecond) whole-brain dynamics in humans (Baillet, 2017, Uhlhaas et al., 2017, Gross, 2019). With millisecond temporal resolution and whole-brain coverage, MEG can bridge the gap between findings obtained by fMRI (which offers excellent spatial resolution with whole-brain coverage, but limited temporal resolution) and invasive electrophysiological methods such as electrocorticography and single-unit recordings (which provide millisecond temporal resolution, but limited brain coverage). In particular, recent advances in MEG analysis using hidden Markov models (HMMs), facilitated in part by the development of parcellation-based approaches for the dimensionality reduction of MEG data (Colclough et al., 2015, Colclough et al., 2016), have revealed the functional roles of rapid whole-brain dynamics in humans (Baker et al., 2014, Quinn et al., 2018, Vidaurre et al., 2018, Fauchon et al., 2022, Rossi et al., 2023). For instance, HMM analysis of MEG data has probed the whole-brain dynamics underlying distinct subprocesses in object perception and working memory (Quinn et al., 2018, Rossi et al., 2023). Thus, MEG combined with HMM offers a promising approach for examining the rapid whole-brain dynamics involved in value-based decision-making processes.

Several MEG studies have investigated the neural mechanisms of value-based decision making (Hunt et al., 2012, Hunt et al., 2013, Azzalini et al., 2021, Hein et al., 2023), although they have focused primarily on individual brain regions rather than whole-brain networks. Hunt et al. (2012) showed that MEG signals in the 2–10 Hz frequency band (roughly corresponding to the delta and theta frequency bands) tracked the subjective values of choice options estimated using a behavioral economic model based on prospect theory. These MEG signals reflected the sum of the option values early in the choice period and their difference later in the choice period. This aligns with the intuition that we first evaluate the value of each option available and then compare the values between options before committing to a choice (Rangel et al., 2008). This study provided significant insights into the long-standing debate over how to interpret value-related fMRI signals, such as whether activation in the ventromedial PFC during decision making represents the value of individual options or the difference in option values. However, because the study focused exclusively on local brain dynamics, it did not address how value-based decision making is related to coordinated 2–10 Hz activity across distributed brain networks.

In the present study, we investigated whether and how rapid whole-brain dynamics in the 2–10 Hz frequency band, evolving over a few hundred milliseconds, are associated with value-based decision making by applying HMM analysis to MEG data. We used a simple binary decision task involving risky and sure options (**Figure 1A**), which has been repeatedly used in previous neuroimaging studies (Levy et al., 2010, Levy and Glimcher, 2011, Tymula et al., 2012, Suzuki et al., 2016). We hypothesized that trial-by-trial whole-brain dynamics derived from HMM analysis, defined by patterns of 2–10 Hz neural activity spanning across distributed brain networks, would track decision variables (such as the subjective values of choice options) estimated by a computational modeling of the participant’s choice behavior. To characterize how value-related processes unfold over time, we analyzed MEG signals time-locked to both the onset of option presentation (option-locked) and the participant’s response (response-locked) (Tanji and Kurata, 1982, Frömer et al., 2024). These analyses showed that the trial-by-trial whole-brain dynamics reflected the subjective value of the risky option (or the choice set as a whole) as well as the difference in subjective values between the options, with distinct temporal profiles unfolding on a subsecond timescale.

**Figure 1.**
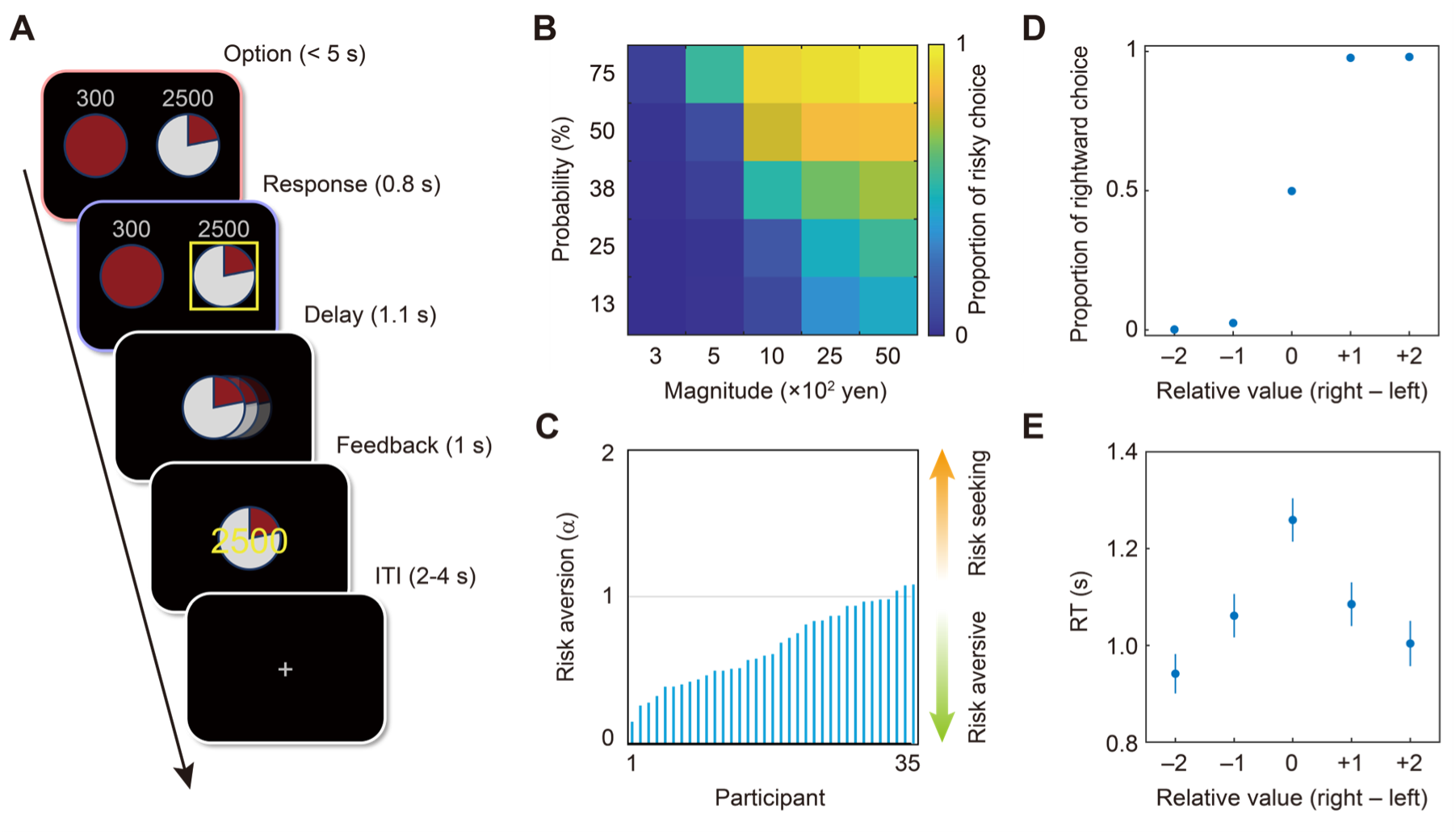
Task and behavioral results. **A** An example of a trial of value-based decision-making task. Participant made a binary choice between a risky and sure option (right/left positions were counterbalanced). Reward magnitude and probability of each option was presented by symbolic numbers and a pie chart, respectively. Upon the participant’s response, the chosen option was highlighted by a yellow frame and moved to the center of the screen. After a delay period, an outcome feedback was presented (won 2500 yen in this example). **B** Proportions of risky choice (averaged across participants) by magnitude and probability of risky option. **C** Risk aversion parameter (a) of individual participants. **D** Proportion of rightward choices as a function of relative value between the right and left options. Relative value was computed by first scaling the trial-wise value difference (right – left) to range from –1 to +1 for each participant and then averaging within each of five equal-interval bins (–1 ≤ x < –0.6, –0.6 ≤ x < –0.2, –0.2 ≤ x ≤ 0.2, 0.2 < x ≤ 0.6, and 0.6 < x ≤ 1, each relabeled as – 2, –1, 0, +1, and +2 for convenience). **E** RT as a function of relative value (right – left). Dots and error bars indicate mean ± SEM across participants (error bars in panel **D** are behind dots; all SEM ≤ 0.0082).

## Results

### Behavioral results

Participants (*N* = 35) underwent MEG measurement while they repeatedly made binary decisions between a risky and sure option (300 trials per participant). The risky option was defined by two attributes, magnitude and probability, which varied from trial to trial. The sure option remained fixed throughout the task (300 yen with 100% probability). Participants responded in 99.20 ± 1.84% (mean ± SD across participants) of the trials within the allotted time (5 s). The mean reaction time (RT) was 1.101 ± 0.233 s (mean ± SD across participants). On average, participants chose the risky option in 40.04 ± 16.41% (mean ± SD across participants) of the trials. In addition, participants chose the risky option more frequently as the magnitude and probability of the risky option increased (**Figure 1B**), indicating that they considered both the magnitude and probability. Using computational models derived from economic theories, we observed that participants’ choice behavior was well described by a model based on prospect theory (variance explained = 0.817 ± 0.069, mean ± SD across participants; see Methods). The risk parameter (α) derived from the model was 0.664 ± 0.266 (mean ± SD across participants), indicating an overall risk-aversive tendency among participants (**Figure 1C**), consistent with numerous previous studies (Kahneman and Tversky, 1979, Levy et al., 2010, Levy and Glimcher, 2011, Tymula et al., 2012, Tymula et al., 2023). Another model-derived parameter (γ) that characterized nonlinearity of the probability weighting function was estimated as 1.046 ± 0.374 (mean ± SD across participants), also consistent with previous research (Tymula et al., 2023). The model-based analysis enabled us to estimate the trial-wise subjective value of each option. As expected, participants chose the option with a higher subjective value in most trials (91.94 ± 5.28%, mean ± SD across participants; **Figure 1D**). Furthermore, the smaller the absolute difference in the values between the two options, the larger the participants’ RTs (**Figure 1E**). That is, participants took longer to make a choice as the values of the two options became more similar. These findings ensured that participants made value-based decisions during the task.

### Whole-brain dynamics associated with visual inputs and motor outputs

We applied HMM to MEG data to examine subsecond whole-brain dynamics. Following the well-established analysis flow used in previous MEG studies (Quinn et al., 2018, Vidaurre et al., 2018, Rossi et al., 2023), we extracted the source-reconstructed and parcellated MEG signals and bandpass-filtered them between the 2 and 10 Hz frequency bands (Hunt et al., 2012). We applied HMM to the parcellated and bandpass-filtered MEG signals to derive a state time series and then examined task-dependent changes in the state time series, referred to as task-evoked occupancy (Quinn et al., 2018). In HMM, the number of states (*K*) is a hyperparameter that must be specified a priori. We chose *K* = 4 in the main analysis, but confirmed that our results were robust across a range of *K* (3–6), as reported in the supplementary figures (**Figures S1–S3**).

First, we examined the task-evoked occupancies that were constant across trials, irrespective of the value of the options, both in option-locked and response-locked manners (**Figure 2**). This demonstrated that the whole-brain patterns of the MEG signal, defined by the combination of parcel-wise means and between-parcel correlations of amplitude envelopes in the 2–10 Hz frequency band (**Figure 2A**), were dynamically modulated by visual inputs and motor execution (**Figure 2B**). For instance, we observed prominent changes in the task-evoked occupancies in the option-locked analysis, peaking at approximately 150–200 ms after the visual presentation of the options (**Figure 2B**). Similarly, as revealed by the response-locked analysis, we observed clear changes in task-evoked occupancies when participants indicated their choice via a motor response (**Figure 2B**). Changes also occurred when the chosen option moved to the center of the screen and when the outcome was presented on the screen (see **Figure 1A** for task events). These findings established that the subsecond whole-brain dynamics captured by the HMM analysis reflected perception- and motor-related changes in the electromagnetic activity of the brain in a temporally resolved manner.

**Figure 2.**
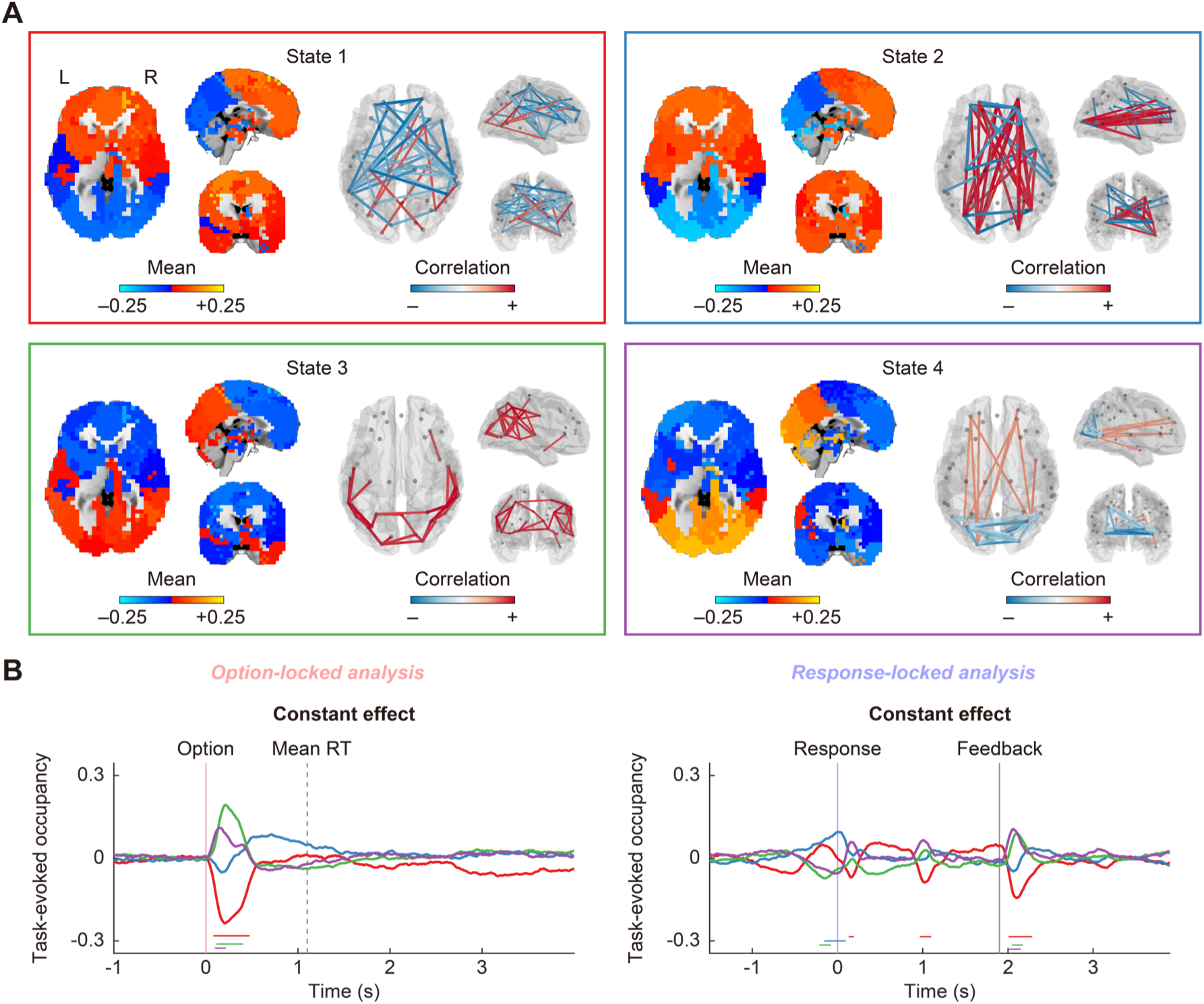
HMM states and task-evoked occupancy. **A** Whole-brain pattern of each HMM state characterized by parcel-wise means and inter-parcel correlations of amplitude envelopes. The mean values are derived from parcel-wise z-scored amplitude envelopes averaged across time points within each state. Correlations (edges) are depicted for parcel pairs exhibiting top 5% strength in absolute value (scaled by maximum absolute value for illustration, which is invariant with respect to 0). R, right hemisphere; L, left hemisphere. **B** Task-evoked occupancy associated with the constant effect across trials. Each time series corresponds to each state shown in Panel A with the same color (red: state 1, blue: state 2, green: state 3, purple: state 4). Horizontal lines at the bottom indicate *P* < 0.05, cluster-based permutation test. The left and right panels show the results of option-locked and response-locked analyses, respectively.

### Whole-brain dynamics associated with value sum and value difference

Next, we asked the critical question of how the task-evoked occupancies were associated with decision variables estimated from the participant’s choice behavior. To this end, we used general linear models (GLMs) to regress HMM state time series onto decision variables on a trial-to-trial basis, as done in previous studies (Quinn et al., 2018; see Methods). Our primary variables of interest were the sum of the subjective values of the two options (*V_chosen_* + *V_unchosen_*, referred to as “value sum”) and the difference in the subjective values between the two options (*V_chosen_* – *V_unchosen_*, referred to as “value difference”), following Hunt et al. (2012). We note that the value sum is mathematically equivalent to the subjective value of the risky option (*V_risky_*) in our task, because the subjective value of the other, sure option (*V_sure_*) was always fixed. Thus, we do not distinguish whether these variables (value sum and *V_risky_*) represent the subjective value of a single option or that of the choice set as a whole. Importantly, we focus on value sum not because we are interested in neural processes related to summation of values, but because this variable can be defined independently of the participant’s choice (Williams et al., 2021, Frömer et al., 2024), affording a straightforward contrast to value difference (*V_chosen_* – *V_unchosen_*). Furthermore, because macroscopic signals measured by MEG pool activity across large neural populations, they are more likely to reflect a mixture of both option values rather than exclusively tracking a single option value (Hunt et al., 2012). The analysis revealed that a specific HMM state (State 1, characterized by a higher value of mean amplitude envelope in the anterior regions of the brain) exhibited changes in task-evoked occupancy in association with these decision variables (**Figure 3AB**). More specifically, in the option-locked analysis, the task-evoked occupancy associated with value sum gradually increased after the option presentation (**Figure 3A**), but plateaued around 700 ms after the option onset. This was accompanied by a decrease in the task-evoked occupancy of another HMM state (State 3, characterized by lower and higher values of mean amplitude envelopes in the anterior and posterior parts of the brain, respectively). We observed a similar pattern of results in the response-locked analysis (**Figure 3A**), with clearer ramping dynamics leading up to the participant’s response and outcome feedback. We also examined whether the observed temporal profiles related to value sum were accounted for by the objective trial attributes (the reward magnitude and probability of the risky option). We found that both reward magnitude and reward probability explained the trial-by-trial variability in the HMM state time series (**Figure S4**), and that the temporal profile of task-evoked occupancies for value sum was more similar to that for reward magnitude than to that for reward probability.

**Figure 3.**
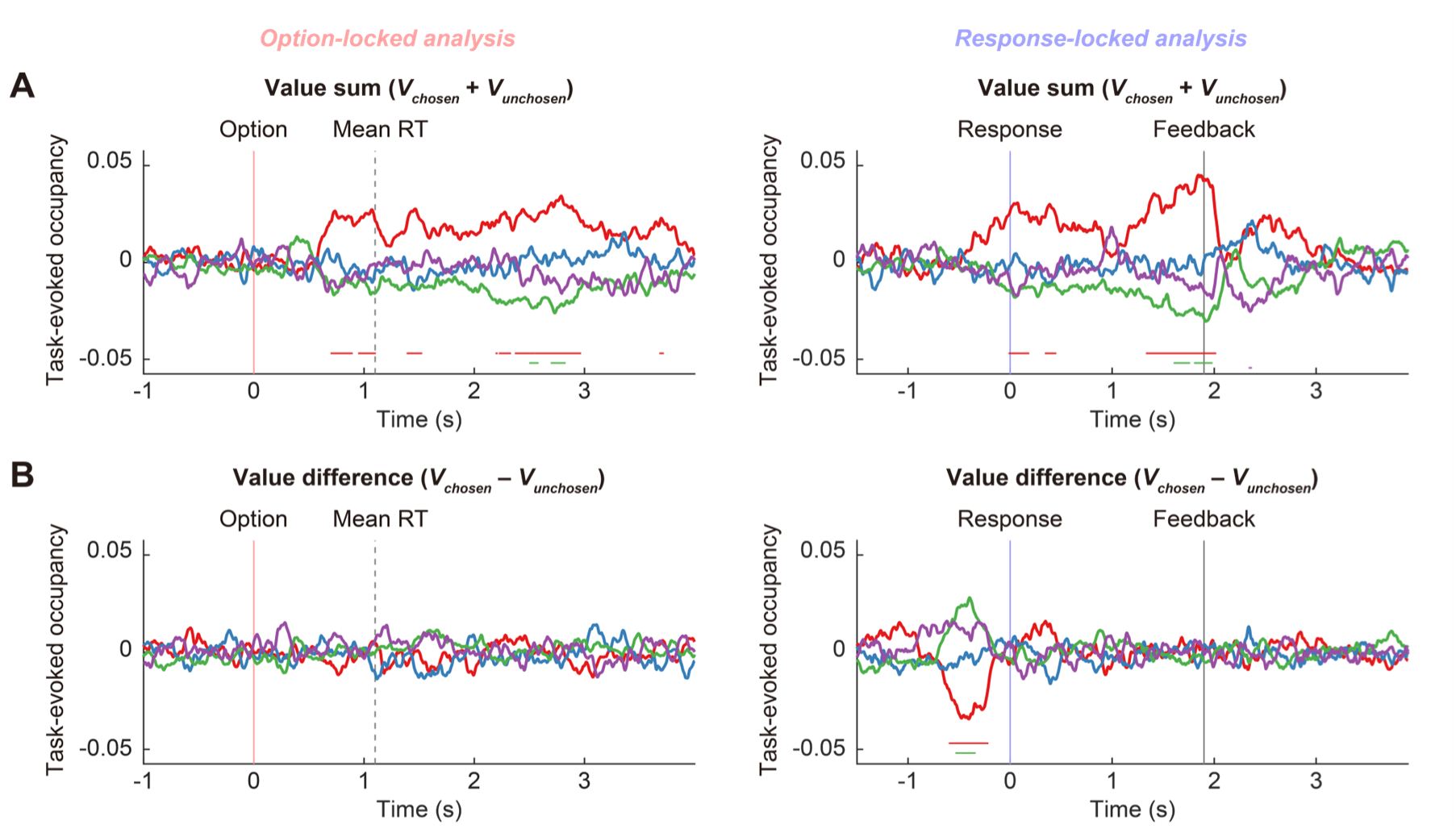
Task-evoked occupancy associated with value sum and value difference. **A** Task-evoked occupancy associated with the value sum (*V_chosen_* + *V_unchosen_*). Note that the value sum is mathematically equivalent to the value of the risky option (*V_risky_*) in our task. **B** Task-evoked occupancy associated with value difference (*V_chosen_* – *V_unchosen_*). The colors of time series correspond to HMM states presented in Figure 2A (red: state 1, blue: state 2, green: state 3, purple: state 4). Horizontal lines at the bottom indicate *P* < 0.05, cluster-based permutation test. The left and right panels show the results of the option-locked and response-locked analyses, respectively.

When we examined the effect of value difference, we found a clear dissociation between the option-locked and response-locked analyses (**Figure 3B**). The option-locked analysis did not show a robust change in the task-evoked occupancy associated with value difference in any HMM state (in other words, the HMM state time series were not modulated by value difference when the time series were time-locked to the option onset). In contrast, the response-locked analysis revealed transient but clear changes in task-evoked occupancies (a decrease in State 1 and an increase in State 3) around – 600 to –200 ms before the participant’s response (**Figure 3B**). To clarify the observed effect of the value difference on the HMM states, we performed two additional GLM analyses in which we modeled the following variables instead of the value difference: i) absolute value difference (|*V_chosen_* – *V_unchosen_*|) or ii) trial-wise RT. The GLM that modeled the absolute value difference yielded virtually the same results as those obtained by the GLM that modeled the value difference (**Figure S5**), which is not surprising given that participants mostly chose the option with a higher value (**Figure 1D**). The GLM that modeled trial-wise RT also yielded similar results (**Figure S5**), reflecting the high correlation between the value difference and RT (median Spearman *r* = –0.342).

To assess the robustness and specificity of our findings, we performed two additional analyses. First, we conducted time-resolved multivariate pattern analysis (MVPA), another approach for probing whole-brain neural dynamics (Liu et al., 2019, Liu et al., 2021, Russek et al., 2024). The MVPA results showed a pattern similar to that obtained with the HMM analysis (**Figure 4**), providing converging evidence for the differential temporal profiles between the option-locked and response-locked analyses in whole-brain dynamics associated with value difference. Second, we performed HMM analyses separately for the 1–4 (delta), 4–8 (theta), 8–13 (alpha), and 15–35 Hz (beta) frequency bands. We observed differential option-locked and response-locked whole-brain dynamics in the 1–4, 4–8, and 8–13 Hz frequency bands, but not in the 15–35 Hz frequency band (**Figures S6–S9**).

**Figure 4.**
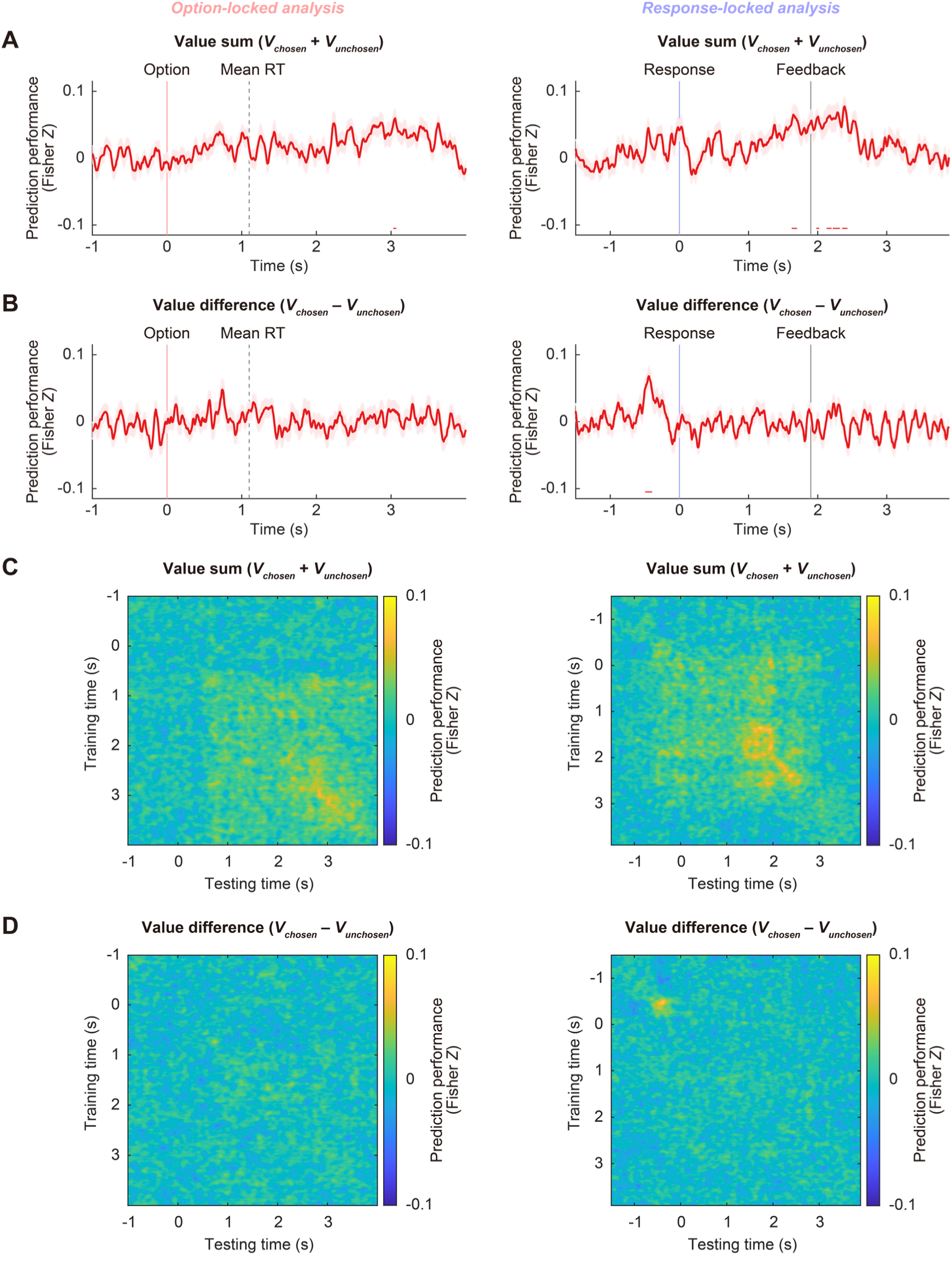
Time-resolved MVPA associated with value sum and value difference. **A** Time-resolved prediction performance (Fisher Z transformed Spearman correlation coefficient) associated with value sum. **B** Time-resolved prediction performance associated with value difference. The line and shaded area in each panel represent the mean ± SE across participants. Horizontal lines at the bottom indicate *P* < 0.05, cluster-based permutation test. **C** Temporal generalization matrix of prediction performance associated with value sum. **D** Temporal generalization matrix of prediction performance associated with value difference. The left and right panels show the results of option-locked and response-locked analyses, respectively.

In summary, we found that trial-by-trial whole-brain dynamics in the 2–10 Hz frequency band were associated with value sum in both the option-locked and response-locked analyses, but were associated with value difference only in the response-locked analysis.

### Whole-brain value dynamics differentiated by participant’s choice

To help uncover how whole-brain value dynamics give rise to choice, we also examined whether the association between trial-by-trial whole-brain dynamics and the value of a choice option depended on the participant’s trial-wise choice (i.e., either choosing the risky or sure option). We reasoned that the trial-by-trial whole-brain dynamics preceding the participant’s response would reflect the value of the risky option (V*_risky_*) to a greater degree in trials in which the participant eventually made risky choices rather than sure choices (Krajbich et al., 2010, Lim et al., 2011). We further reasoned that the trial-by-trial whole-brain dynamics during the anticipatory period (defined as the period after the participant’s response and before outcome feedback) would be associated with V*_risky_* only when the participant had chosen the risky option. To formally test this idea, we defined a binary variable denoting the participant’s trial-wise choice (referred to as “choice type”; risky = +1, sure = –1), and examined whether choice type modulated the effect of V*_risky_* on the HMM time series. This analysis revealed an interaction effect between V*_risky_* and choice type in the response-locked analysis, observed approximately –550 to –250 ms before and 900–950 ms after the participant’s response (**Figure S10A**). A post-hoc analysis clarified that this interaction effect was predominantly driven by temporal changes in the task-evoked occupancy associated with V*_risky_* in the risky-choice trials (**Figure S10B**), but not in the sure-choice trials (**Figure S10C**). To determine the possible origin of this interaction effect, we performed two additional GLM analyses, in which we modeled the expected value (*EV_risky_*) or variance (*VAR_risky_*) of the risky option instead of V*_risky_*. This follows standard frameworks in economics and finance, in which a risky asset (defined in terms of a probability distribution) is characterized by its expected value and variance (Suzuki et al., 2016, Williams et al., 2021). We found an interaction effect similar to that described above for *EV_risky_*, but not for *VAR_risky_* (**Figure S11**). In summary, whether the participant chose the risky or sure option differentiated the trial-by-trial whole-brain dynamics associated with the value of the risky option, which was primarily driven by the expected value of the risky option in the risky-choice trials.

### Whole-brain dynamics associated with outcome feedback

Lastly, we investigated whether whole-brain dynamics were modulated by outcome feedback. We ran GLMs that included either i) the subjective value of the outcome (*V_feedback_*) or ii) reward prediction error (PE) as a parametric regressor. We found that *V_feedback_* was associated with trial-by-trial whole-brain dynamics, specifically State 1, approximately 900–950 ms after feedback onset (**Figure S12A**). We also found that PE was associated with trial-by-trial whole-brain dynamics (State 4) approximately 100–150 ms after feedback onset (**Figure S12B**). Given the short latency, the early PE-related effect may have partly reflected the perceptual difference between the win and loss outcomes (visually presented as symbolic numbers, with three to four digits [300–5000 yen] in the win outcomes and one digit [0 yen] in the loss outcome). We further observed a noticeable change in the task-evoked occupancy associated with PE approximately 900 ms after feedback onset, although this effect was not significant (**Figure S12B**). In exploratory HMM analyses targeting different frequency bands, we found that trial-by-trial whole-brain dynamics were associated with PE in each of the 1–4, 4–8, 8–13, and 15–35 Hz frequency bands, with temporal profiles varying across frequencies (**Figure S13**). Overall, these results indicate that the trial-by-trial whole-brain dynamics associated with decision variables during the choice and anticipatory periods are also related to value processing after outcome feedback.

### Local brain dynamics

Our central investigation is regarding whole-brain dynamics derived from the HMM analysis. However, we performed an additional conventional analysis focusing on local brain activity to verify that our MEG signals capture simple brain responses related to visual and motor processing. Specifically, we performed a source-level GLM analysis on a 5 mm grid space (as opposed to the parcellated time series used in the HMM analysis). We observed clear visual responses in the primary visual cortex at the timings of visual stimulations (option presentation, highlighting the chosen option, and outcome feedback) and motor responses in the primary motor cortex at the time of the participant’s response (**Figure 5**).

**Figure 5.**
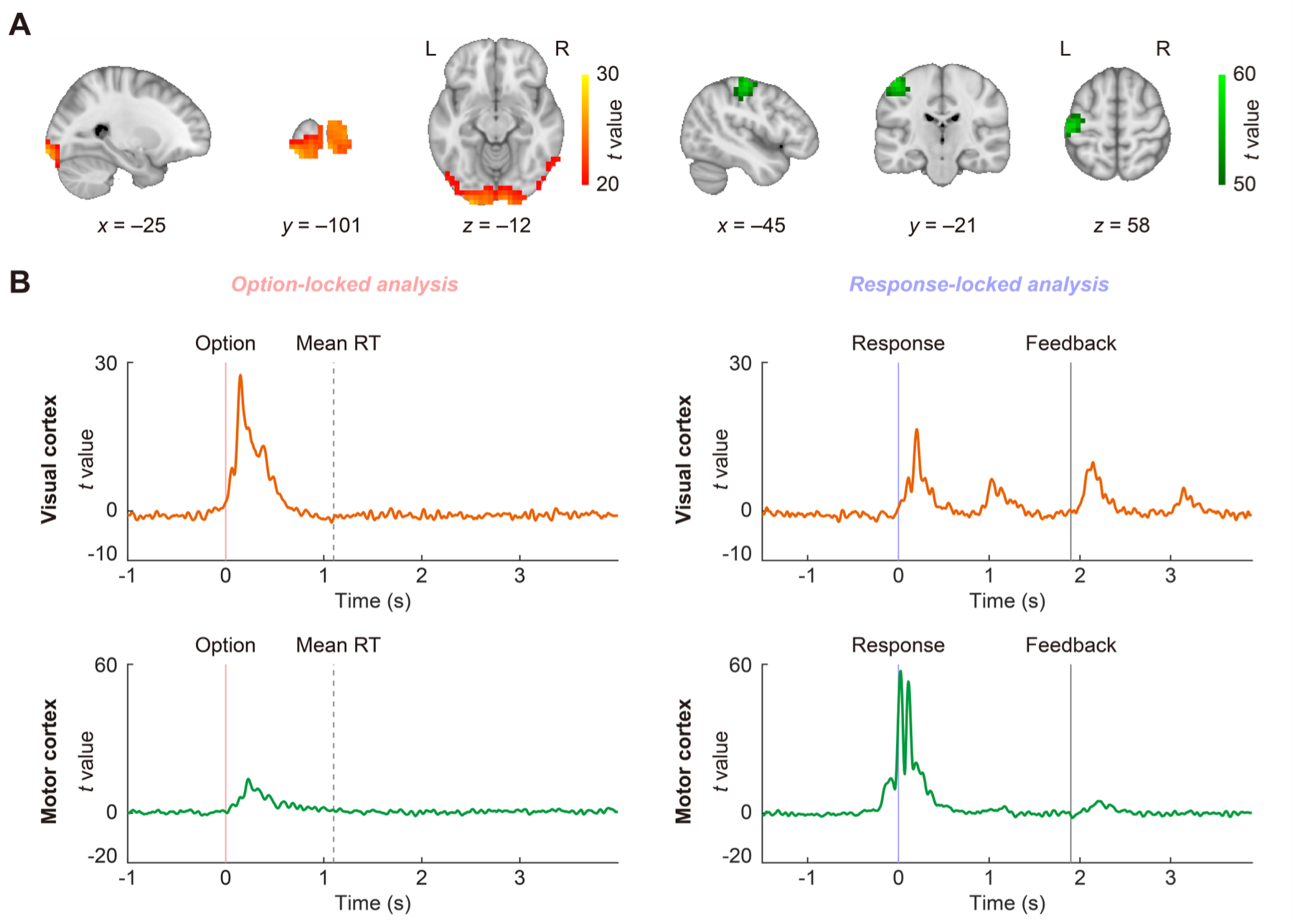
Source-reconstructed activation associated with perceptual inputs and motor execution. **A** Group-level activation maps derived from source-level GLMs assessing the constant effect across trials (left: option-locked analysis, right: response-locked analysis). Colors indicate grid-wise maximum *t* values across time points (maximum intensity projection). MNI coordinates indicate the global peak (across time points and grids) identified in each GLM. R, right hemisphere; L, left hemisphere. **B** Group-level time series of *t* value (estimated at each time point) at the peak grids shown in panel A (visual cortex: –25, –101, –12; motor cortex: –45, –21, 58).

In addition, we performed a time-frequency analysis in a set of regions of interest (ROIs) selected a priori. In this analysis, we expanded our frequencies of interest to 1–45 Hz for an exploratory purpose (Quinn et al., 2018). The results revealed robust and widespread suppression in the beta frequency band (15–35 Hz, “beta suppression”) around the time of the participant’s motor execution and subsequent rebound (“beta rebound”), which are hallmarks of the electromagnetic brain responses related to motor execution (Cheyne, 2013). We also observed widespread task-related desynchronization in the alpha frequency band (8–13 Hz, “alpha suppression”), which is typically observed in association with various cognitive processes, such as attention (Hari and Salmelin, 1997). We also report the results of ROI-wise GLM analyses associated with value sum and value difference (**Figures S14 and S15**). These results shed light on regionally and spectrally structured patterns of the effects of value processing on local brain dynamics; however, these are only for descriptive purposes and are not subject to statistical significance testing.

## Discussion

In this study, we aimed to elucidate whole-brain neural dynamics underlying value-based decision making in humans. First, we applied HMM analysis to MEG signals (2–10 Hz frequency band) and obtained a concise set of whole-brain dynamics in a data-driven manner (without references to task information such as stimulus/response timings and trial attributes). Then, we asked whether any of these whole-brain dynamics (the HMM state time series) were associated with trial-wise decision variables derived from a behavioral economic modeling of participants’ choices. As a result, we identified subsecond whole-brain dynamics tracking several aspects of the value-based decision-making process. The temporal profiles of the dynamics likely reflected perceptual processing of the options presented, subjective valuation of the options (or the choice set), preparation and execution of action (implementation of a choice), as well as anticipatory processing until the delivery of the outcome feedback (summarized in **Figure 6**). Importantly, we found a dissociation of the patterns of results between the analysis time-locked to the option onset and that time-locked to the participant’s response, which provides insights into whether the observed relations between whole-brain dynamics and decision variables (value sum and value difference) were more linked to processing of perceptual inputs or to motor outputs. Furthermore, we observed two types of subsecond whole-brain ramping dynamics, one leading to the participant’s choice and the other to outcome feedback (**Figure 6**). Our findings highlight the critical relevance of whole-brain patterns of oscillatory neural activity (particularly in the 2–10 Hz frequency band) in value-based decision-making processes in humans.

**Figure 6.**
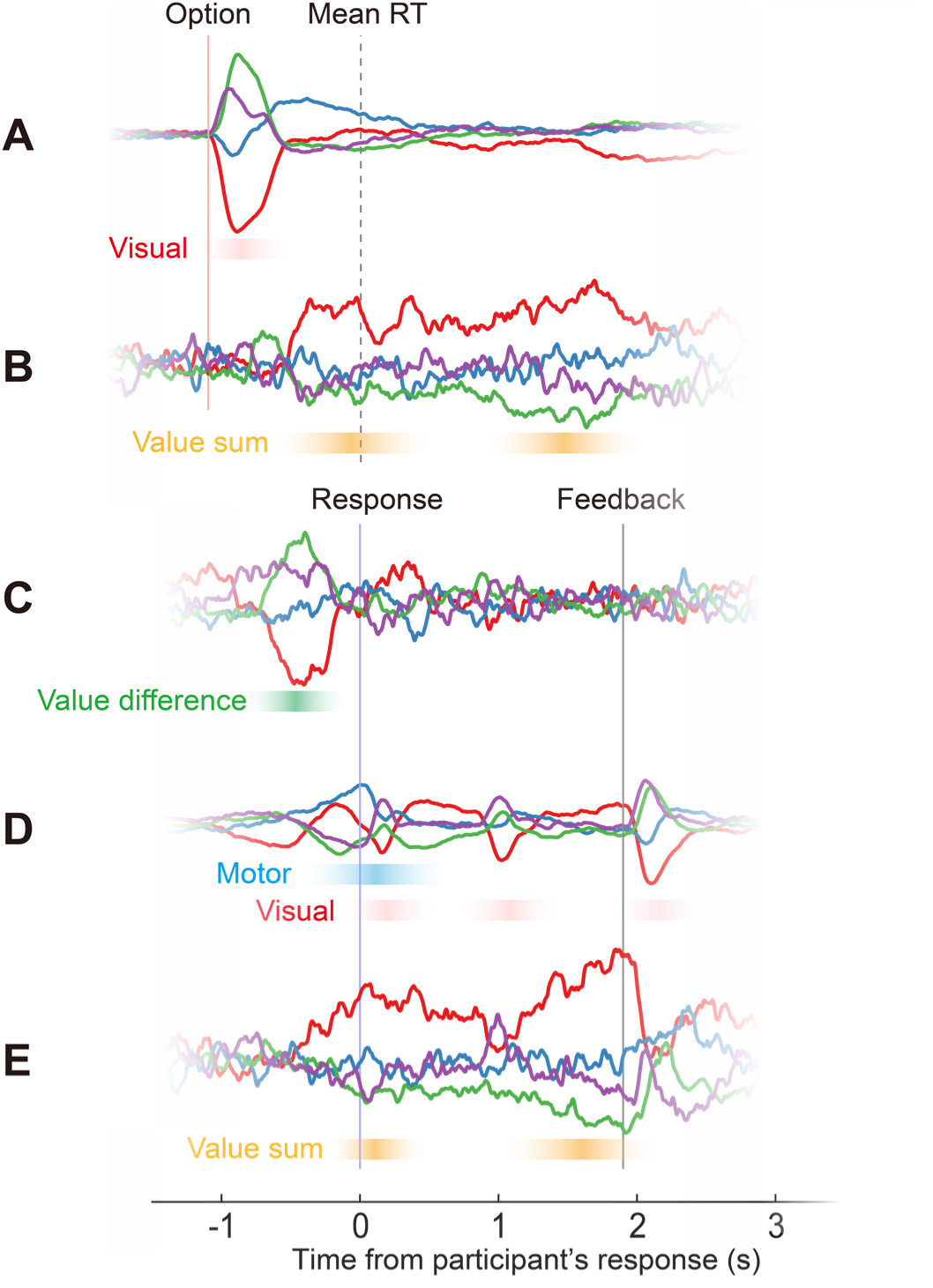
Summary of key findings. **A** Task-evoked occupancy associated with constant effect in option-locked analysis (from Figure 2B). **B** Task-evoked occupancy associated with value sum in option-locked analysis (from Figure 3A). **C** Task-evoked occupancy associated with value difference in response-locked analysis (from Figure 3B). **D** Task-evoked occupancy associated with constant effect in response-locked analysis (from Figure 2B). **E** Task-evoked occupancy associated with value sum in response-locked analysis (from Figure 3A).

Our study demonstrates that HMM analysis of MEG data can effectively capture whole-brain dynamics underlying value-based decision making. Our findings extend the recent lines of MEG research that have dramatically advanced our understanding of the rapid neural mechanisms involved in value-related processes, such as the functional roles of sequential replay in reward, memory, and learning (Kurth-Nelson et al., 2015, Kurth-Nelson et al., 2016, Liu et al., 2019, Liu et al., 2021, Wise et al., 2021, McFadyen et al., 2023, Wimmer et al., 2023). Notably, previous neuroimaging studies (such as fMRI) have often focused on the representation of certain decision variables in individual brain regions (Huettel et al., 2005, Levy et al., 2010). However, recent studies have emphasized value-related computations instantiated by distributed neural networks spanning across the whole brain (Vickery et al., 2011, Hunt and Hayden, 2017). Our results clearly indicate that non-invasively measured whole-brain dynamics at the macroscopic level do reflect value-based decision-making processes unfolding over time.

We employed a binary value-based decision-making task that involved economic risk (Levy and Glimcher, 2011), a standard task in neuroimaging and electrophysiological research for humans and animals (Hunt et al., 2012, Tymula et al., 2012, Hunt et al., 2013, Suzuki et al., 2016, Williams et al., 2021, Imaizumi et al., 2022, Tymula et al., 2023). A previous MEG study using a similar task showed that, when focusing on option-locked activity in the 2–10 Hz frequency band, the MEG signals in the vmPFC and intraparietal sulcus first covaried with the value sum and then with the value difference (Hunt et al., 2012). In our HMM-based analysis using whole-brain activity patterns, we did not find the same pattern in the option-locked analysis. This might be because the participants’ trial-wise RTs spread out in time to a great degree in our study (∼0.5 to 5 s; participants were allowed to make a choice for up to 5 s), which would make a transient, response-locked effect less aligned with the option onset and difficult to detect using option-locked analyses. In addition, HMM-based analyses may be less sensitive to temporal characteristics that are unique to a single or a few local brain regions. Rather, we found another type of temporal dissociation: the effects of value sum on whole-brain dynamics (the HMM state time series) were observed in the option- and response-locked analyses, whereas the effects of value difference were observed only in the response-locked analysis. The dissociable temporal characteristics (value sum vs. value difference) are manifested in different ways in the study by Hunt et al. (2012) and our study; however, they likely stem from the same temporal evolution of decision-making processes in the brain. That is, neural correlates of value sum arise as participants evaluate a choice set (each option or the set as a whole) before committing to a specific choice (thus, option-locked), whereas neural correlates of value difference emerge as participants increasingly lean toward one option over others and eventually reach a decision (thus, response-locked).

With a closer look, the effect of value difference on the response-locked HMM state time series was observed around –600 to –200 ms with respect to the participants’ response (**Figure 3B**). Given that the average reaction time was around 1.1 s, the observed dynamics associated with value difference emerged substantially after the option onset, suggesting that many internal operations took place in the brain before value difference was robustly manifested in whole-brain dynamics. In fact, the simple sensory/perceptual effects of option presentation (irrespective of the option value) on the HMM state time series were observed around 100–500 ms after option onset (**Figure 2B**), which is clearly distinct from the effect of value difference. It is also clear that the whole-brain dynamics associated with value difference are distinct from the dynamics associated with simple (value-irrelevant) motor responses primarily observed around –200 to 0 ms before motor response (**Figure 2B**). These observations confirm that the relationship between value difference and whole-brain dynamics cannot be attributed to simple visual- or motor-related processes.

The functional role of neural correlates of value difference is currently under debate. There are at least two possible accounts. One account argues that neural correlates of value difference directly reflect value comparison processes, often assuming neural circuit models in which two populations of neurons (in binary decision cases) encode the value of each option and their activity change over time via mutual inhibition (Wunderlich et al., 2009, Hare et al., 2011, Hunt et al., 2012, Hunt and Hayden, 2017). Another account argues that neural correlates of value difference are not directly responsible for value comparison per se, but reflect associated processes, such as attention and motivation (Shenhav et al., 2016, Frömer et al., 2024). For instance, when the values of two options are similar (a more difficult choice), the brain recruits attentional resources to a greater extent so that the decision maker can scrutinize the value of each option to make a better choice (Frömer et al., 2024). Notably, the HMM state that was most robustly associated with value-based decision making in our results (State 1, whose task-evoked occupancy increased with the value sum) *decreased* its occupancy as a function of value difference (**Figure 3B**). In other words, the brain expressed this HMM state more frequently when the values of the two options were similar. This suggests that an increase in the occupancy of this HMM state cannot simply be interpreted as reflecting positive affect associated with greater option value (“pleasantness” or “excitement”). Rather, the result of our GLM analysis that modeled trial-wise RT suggests a possible link to choice difficulty. However, no single interpretation would fully characterize the true nature of this HMM state; the temporal changes in this HMM state time series (either increase or decrease) should be viewed as a macroscopic manifestation of underlying value-related computational processes, to which past research has given many different (but interrelated) interpretations, such as pleasant or anxious feelings, heightened attention or motivation, and the need for resolving uncertainty or cognitive control, amongst others (Shenhav et al., 2013, Hunt and Hayden, 2017).

Several studies have explicitly tested which of the value comparator or choice difficulty account better explains brain activity in certain brain regions (such as the pre-supplementary motor area [preSMA]), for instance by employing classes of neural circuit or drift diffusion models (DDM) that predict different patterns of neural responses between “correct” (participant choosing the option with a greater subjective value) and “error” (choosing the option with a smaller subjective value despite availability of a better-valued option) trials (Wunderlich et al., 2009, Hare et al., 2011). However, distinguishing between these two accounts is very challenging. In fact, some studies have pointed out that brain activity whose patterns closely followed theoretical predictions of the value comparator model could actually be better explained by choice difficulty (Shenhav et al., 2014, Shenhav et al., 2016). In the current study, we could not adopt the strategy exploiting the different responses between the “correct” and “error” trials because the “error” trials were rare (**Figure 1D**). In addition, whether the temporal evolution of our signals of interest (HMM state time series) can be adequately captured by typical value comparator models such as DDMs is not warranted (Frömer et al., 2024). Thus, the exact nature of our findings regarding whole-brain dynamics associated with value difference remains to be elucidated in future studies.

A particularly intriguing finding in our study is the two ramping dynamics: one leading to the participant’s response and the other leading to the outcome feedback. These two ramping dynamics were observed in the same HMM state. Hence, it is possible that both dynamics, at least in part, reflected the common aspects of value processing, such as representing uncertainty related to action and outcome values (Bach and Dolan, 2012, Soltani and Izquierdo, 2019, Matsumoto et al., 2022). However, the differences between the two dynamics were also notable. The ramping dynamics before choice may mirror the evidence accumulation process assumed in DDMs (Polanía et al., 2014, Pisauro et al., 2017), although a ramping-up temporal profile does not necessarily mean that it directly reflects evidence accumulation (as discussed above in relation to the value comparator vs. choice difficulty accounts). The other ramping dynamics before outcome feedback cannot be explained by evidence accumulation (nor by action selection or motor preparation) because no choice or action takes place at this stage of the trial. Rather, these dynamics may be related to dopamine functions associated with upcoming rewards (Lerner et al., 2021, Gershman et al., 2024). Animal studies have shown that the activity of dopamine neurons ramps up as an animal approaches (spatially or temporally) rewards (Kim et al., 2020, Sosa and Giocomo, 2021). Such a function may be crucial to acquire a value map of the environment and to guide adaptive behavior. Similar brain mechanisms may also play roles in humans to signal the expected timing and value of upcoming rewards. However, caution is required because similar temporal profiles (ramping up) could emerge from other mechanisms not directly related to dopamine. Experiments using pharmacological manipulation (such as acute phenylalanine/tyrosine depletion before task-MEG recordings) may allow us to empirically test whether the dopamine system is involved in the ramping dynamics observed in the current study. Another line of studies relevant to ramping dynamics before outcome feedback is grounded in economic theories on anticipatory utility (Loewenstein and Lerner, 2003, Loewenstein, 2006). A few fMRI studies using temporal discounting tasks have examined the neural correlates of anticipatory utility dynamics and reported that BOLD signals (for instance, in the anterior prefrontal cortex) ramped up until the delivery of temporally delayed rewards (Jimura et al., 2013, Iigaya et al., 2020, Tanaka et al., 2020). These studies typically used a delay period of ∼30–60 s to probe the temporal profiles of BOLD signal dynamics. However, the limited temporal resolution of fMRI and the sluggish nature of BOLD signals hinder the clarification of the subsecond neural dynamics involved in value processing (Shidara and Richmond, 2002, Aquino et al., 2023, Balewski et al., 2023, Man et al., 2024). Our frequency-specific HMM analysis provides further insights into these ramping dynamics (**Figures S6–S9**). In the 1–4 Hz (delta) and 4–8 Hz (theta) frequency bands, the overall temporal characteristics of the task-evoked occupancies resembled those in the 2–10 Hz frequency band, as expected given their overlapping frequency ranges. In contrast, in the 8–13 Hz (alpha) and 15–35 Hz (beta) frequency bands, we observed an additional HMM state that exhibited ramping dynamics only leading up to the outcome feedback but not at the time of the participant’s response (**Figures S8 and S9**). These frequency-specific ramping dynamics associated with value processing would have been difficult to detect using fMRI. MEG may offer unique opportunities to unveil subsecond whole-brain dynamics associated with time-discounted values, anticipatory processes, and other types of dynamically changing values (Janssen and Shadlen, 2005, McGuire and Kable, 2015). Elucidating the neural mechanisms underlying these dynamic value computations may open new avenues for understanding maladaptive decision making in clinical populations, such as patients with obsessive-compulsive and gambling disorders (Uhlhaas et al., 2017, Uhlhaas et al., 2018, Durstewitz et al., 2021, Huys et al., 2021, Suzuki et al., 2023).

This study has several limitations. First, although we used a well-established value-based decision making task (Levy and Glimcher, 2011), it involved only two trial-to-trial attributes (magnitude and probability of one option) and was not optimal for decorrelating interdependent decision variables. For instance, we did not distinguish value sum from the value of the risky option because participants always chose between a risky option and a fixed reference option. This was an inevitable consequence of our design consideration of prioritizing simplicity of the task. Future studies could incorporate additional trial-to-trial attributes to more clearly disentangle interdependent variables related to valuation and choice. Second, we had an a priori focus on the 2–10 Hz frequency band, capitalizing on the previous MEG study revealing a critical involvement of this frequency band in value-based decision making (Hunt et al., 2012). However, other frequency bands may also play key roles. For instance, reward processing subserved by the dopamine system (including the front-striatal network) has been linked to beta oscillations (Schwerdt et al., 2020, Hoy et al., 2024). Indeed, we found an HMM state time series associated with reward prediction error (PE) in the beta frequency band (**Figure S13**). Intriguingly, however, PE-associated HMM dynamics were also observed in the delta, theta, and alpha frequency bands, implying that reward prediction error affects whole-brain dynamics over a broad frequency spectrum. In addition, invasive electrophysiological recordings from single neurons in humans have shown that certain decision variables (value of the chosen option) are correlated from trial to trial with spike activity in local brain regions (Aquino et al., 2023), which is thought to be tied to high-gamma oscillations (Jensen et al., 2007). Further research is needed to clarify the functional roles of oscillatory neural activity in various frequency bands and its inter-frequency coupling. Third, we focused on research questions regarding whole-brain dynamics and did not attempt to relate individual brain regions to specific decision variables. We emphasize that this approach aligns with the view that decision making emerges from the dynamics of the brain as a distributed system and cannot be fully understood by examining individual brain regions in isolation (Hunt and Hayden, 2017). Nonetheless, our supplementary ROI analysis (**Figures S14 and S15**) did provide some hints on functional specificity in individual brain regions (although we strictly avoided reverse inference). These results may help generate hypotheses about locally specialized brain functions related to value processing, which should be tested in independent datasets obtained in future studies.

Despite these limitations, our study demonstrates the advantages of combining MEG recordings with HMM analyses to elucidate subsecond whole-brain dynamics that support value-based decision making in humans. To our knowledge, this study provides the first evidence that 2–10 Hz whole-brain dynamics identified by HMM analysis reflect trial-wise subjective value derived from a formal economic theory (i.e., prospect theory). The approach used here may help bridge findings from fMRI research (often limited in temporal resolution) and human single-unit recordings (often limited in brain coverage), as well as translate results from large-scale simultaneous neural recordings in animals to human brain mechanisms (Urai et al., 2022, Stringer and Pachitariu, 2024). Research highlighting the relevance of fast-evolving whole-brain dynamics in value processing, like ours, may facilitate integration of knowledge across modalities and species and contribute to understanding how value and choice arise from distributed neural dynamics.

## Methods

### Participants

We recruited 40 healthy adults with normal or corrected-to-normal vision and no self-reported history of psychiatric or neurological disorders. We excluded five participants from the analysis because of the following reasons: inability to achieve vision correction with non-metallic glasses (*N* = 1), falling asleep and not responding in ∼29% of trials (*N* = 1), missing trigger information due to technical issues (*N* = 1), and imprecise coregistration between MEG and MRI data (*N* = 2). The remaining 35 participants (13 females, age: range = 20–44 years, mean = 26.2 years, SD = 7.1 years) were included in the behavioral and MEG analyses. All participants provided written informed consent before the experiment, and the Ethics Committee of the National Center of Neurology and Psychiatry approved this study.

### Experimental task

Participants performed a gambling task while they undertook MEG measurement (**Figure 1A**). In each trial, the participant made a binary decision between a sure option and a risky option. Each option was defined by combining a reward magnitude and a reward probability. The reward magnitude and probability of the risky option varied from trial to trial. The magnitude was selected from 300, 500, 1000, 2500, or 5000 yen, and the probability from 13, 22, 38, 50, or 75%, yielding 25 possible combinations of magnitude and probability for the risky option (**Figure 1B**). The sure option always had a fixed reward magnitude of 300 yen and a reward probability of 100%. At the beginning of each trial, the risky and sure options were displayed on the screen as a pair of numbers accompanied by pie charts. The reward magnitudes were indicated by numbers above the pie charts, and the reward probabilities were indicated by the areas colored red in the pie charts. The options remained on the screen until the participant made a response (by right index or middle fingers) or up to 5 s. After the participant’s response, the chosen option was highlighted by a yellow rectangle for 0.8 s. Subsequently, the unchosen option disappeared, and the chosen option moved to the center of the screen over 0.1 s and remained there for 1 s. This was followed by a 1-s presentation of outcome feedback (displayed as a yellow number): If the participant had chosen the risky option, the outcome was either a win (obtaining a magnitude of reward stated by the option) or loss (obtaining 0 yen); if she or he had chosen the sure option, the outcome was 300 yen. After an inter-trial interval (ITI) of 2–4 s, the next trial began. Participants performed 300 trials divided into four runs (75 trials each lasting approximately 10 min). The positions of the two options (the right or left side of the screen) were fixed within a run and counterbalanced across runs, and the position of the risky option (right/left) in the initial run was randomized across participants. At the end of the experiment, one trial was randomly selected, and its outcome was added as a bonus to the base payment of 5000 yen (or 7000 yen for the participants who took part in the structural MRI scan on the same day).

### Behavioral data analysis

We performed a model-based analysis to derive subjective value from the participants’ choice behavior. First, the subjective value (*V*) of an option was modeled by a utility function based on the prospect theory (Kahneman and Tversky, 1979, Hunt et al., 2012, Tymula et al., 2023):

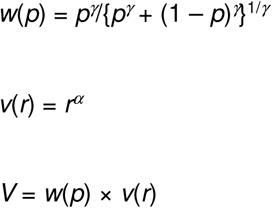

where *r* and *p* are the reward magnitude and probability, respectively, and *w*(*p*) is a probability weighting function parameterized by *γ*. Here, *α* = 1 indicates risk-neutral behavior, whereas *α* < 1 and *α* > 1 indicate risk-aversive and risk-seeking behaviors, respectively. In our binary choice setting, the probability of choosing Option A (*P_A_*) over Option B was modeled by a softmax function:

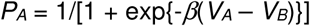

where *V_A_* and *V_B_* indicate the subjective values of Options A and B, respectively. We estimated the free parameters (*α*, *β*, and *γ*) separately for individual participants by minimizing negative log likelihood using *fmincon* in MATLAB R2024a, with the following constraints on the parameters: *α* = [0 2], *β* = [0 Inf], *γ* = [0 2]. These constraints were determined based on the typical ranges of these parameters reported in previous studies (such as Tymula et al., 2023).

We also considered two alternative models in which the subjective value of an option is represented by different utility functions. The first one was an exponential utility function (Levy and Glimcher, 2011, Suzuki et al., 2016, Williams et al., 2021):

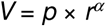

The second one was a mean-variance utility function (Suzuki et al., 2016, Williams et al., 2021):

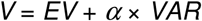

where *EV* and *VAR* indicate the expected value (*r* × *p*) and variance (*r*^2^ × *p* × (1 – *p*)), respectively.

The choice probability was modeled by a softmax function in the same manner as the primary model. The model fit was evaluated using Bayesian Information Criteria (BIC) computed for each participant. The across-participant summation of BIC was 4779.87 for the prospect theory model (PT), 4826.02 for the exponential utility model (EU), 6272.47 for the mean-variance utility model (MV), showing that PT had the best fit. The risk-aversion parameters estimated by these models were highly correlated with each other across participants (Spearman *r* = 0.959 between PT and EU, *r* = 0.946 between PT and MV, and *r* = 0.992 between EU and MV; **Figure S16**). In addition, all these risk-aversion parameters were highly correlated with the proportion of risky choices, which is a model-independent measure of individual risk attitudes (Spearman *r* = 0.933 for PT, *r* = 0.961 for EU, and *r* = 0.945 for MV; **Figure S16**).

We examined the relationship between the model-derived subjective value with the participant’s trial-wise choice and RT as follows (**Figure 1D and E**). First, we computed the “relative value” (right – left), which was obtained by scaling the trial-wise value difference between the right and left options (*V_right_* – *V_left_*) to range from –1 to +1 for each participant. Next, we defined five equal-interval bins (–1 ≤ x < –0.6, –0.6 ≤ x < –0.2, –0.2 ≤ x ≤ 0.2, 0.2 < x ≤ 0.6, and 0.6 < x ≤ 1, each relabeled as –2, –1, 0, +1, and +2 for convenience), and calculated the proportion of risky choices and mean RT within each bin. The resulting bin-wise proportion of risky choices and mean RT were averaged across participants.

### MEG and MRI data acquisitions

MEG data were recorded in a sitting position using a 306-channel MEG system (Elekta Neuromag TRIUX; MEGIN Oy, Croton Healthcare, Helsinki, Finland). The data were recorded in a magnetically shielded room at the National Center of Neurology and Psychiatry. MEG signals were sampled at 1,000 Hz and bandpass-filtered online between 0.1 and 330 Hz. Five head-position indicator (HPI) coils were used to continuously track the participants’ head motions. The positions of the HPI coils relative to the participant’s head were measured using a three-dimensional digitizer (Polhemus Isotrack) before MEG measurement. Vertical electrooculograms (EOG) and electrocardiograms (ECG) were recorded to detect eyeblinks and heartbeats, respectively.

MRI data were acquired using a 3T MRI scanner equipped with a 32-channel head coil (Siemens Skyra; Siemens Healthcare, Erlangen, Germany). An anatomical T1-weighted image with a magnetization-prepared rapid gradient echo (MPRAGE) sequence (repetition time = 2500 ms, echo time = 2.18 ms, flip angle = 8°, resolution = 0.8 mm × 0.8 mm × 0.8 mm) was obtained for each participant.

### MEG data analysis

#### MEG data preprocessing

After denoising with Sig3-Denoise (Larson and Taulu, 2017) and signal-space separation with MaxFilter (version 2.2; Taulu et al., 2006), the MEG data were preprocessed using the Oxford Centre for Human Brain Activity (OHBA) Software Library (OSL) run on MATLAB R2024a. First, MEG data were coregistered with the participant-specific T1-weighted structural MR image using the RHINO algorithm, high-pass filtered at 0.5 Hz, and downsampled to 200 Hz. Artifact rejection was performed using the AFRICA algorithm, whereby independent components that correlated with the ECG and/or EOG (r > 0.5) were removed. Bad segments were marked by dividing the continuous MEG time series into 1-s non-overlapping windows and then detecting the windows with outliers, identified based on the generalized extreme Studentized deviate method (Rosner, 1983) implemented in OSL. These bad segments were excluded from subsequent analyses.

#### Source reconstruction

To obtain MEG signals in the source space, we performed source reconstruction as previously reported (Quinn et al., 2018, Rossi et al., 2023). First, we normalized the MEG sensor data across sensor types (magnetometers and gradiometers) using an eigenvalues decomposition algorithm (Woolrich et al., 2011). Next, we constructed a single-shell forward model and applied a beamforming algorithm (the bilateral beamformer, Woolrich et al., 2011) with covariance computed within a 2–10 Hz frequency band to project the sensor data to the source space. We used a 5-mm grid in the MNI space.

#### HMM

We performed HMM analysis, following a previous study that applied the HMM analysis to MEG data (Baker et al., 2014, Quinn et al., 2018, Vidaurre et al., 2018, Fauchon et al., 2022, Rossi et al., 2023). Using a weighted (nonbinary) parcellation with 39 brain regions, which has been frequently used in previous MEG research (Colclough et al., 2016, Quinn et al., 2018, Kohl et al., 2024), the source-reconstructed MEG data were parcellated to obtain a parcel-wise MEG time series. We extracted the time series of each parcel as the first principal component across the time series of the voxels included in the parcel. To attenuate the spatial leakage of source-space MEG signals (correlations of time series between neighboring regions), we applied the symmetric multivariate leakage correction method (Colclough et al., 2015). Next, we computed the amplitude envelope (2–10 Hz) of the parcel-wise MEG time series using the Hilbert transform. The envelopes were smoothed with a 50-ms (10 time points) moving average filter and z-score normalized for each parcel. The periods marked as bad segments were discarded, and the remaining continuous “good segments” were concatenated (with the information about discontinuity in the time series considered in the following HMM estimation step). The resulting time series was further concatenated across participants and sessions, and subjected to HMM inference (that is, the HMM inference was performed at the group level). HMM was estimated using the HMM-MAR toolbox (Quinn et al., 2018). We used variational Bayes inference implemented in the HMM-MAR toolbox, adopting the stochastic inference option with the same parameters used in Quinn et al. (2018) (maximum number of cycles = 500, tolerance = 10^-7^, forget rage = 0.7). We estimated the state time series (*T* × *K*, where *T* is the total number of time points included in the HMM analysis and *K* is the number of state), *M* × 1 vector *μ_k_* indicating parcel-wise means (where *M* is the number of parcels) and *M* × *M* covariance matrix *Σ_k_* representing the statistical dependence (“functional connectivity”) of envelope time series between parcel pairs for each state. The model assumed that *μ_k_* and *Σ_k_* follow multivariate normal distribution. We visualized *μ_k_* as a spatial map (for each state) indicating the parcel-wise means of the amplitude envelopes, and we also visualized power correlation between parcel pairs (**Figure 2A**). We repeated the HMM inference 10 times and determined the estimation result that gave the lowest free energy for a given number of states (Quinn et al., 2018). We varied the number of states (*K* = 3–6) and reported the results with *K* = 4 in the main text. We confirmed that our main findings regarding the HMM analysis were robust to the selection of the number of states (similar patterns of states were observed with other *K*; see **Figures S1–S3**).

#### GLM

To examine how the HMM state time series was associated with decision variables (value sum and value difference), we performed GLM analyses in which epoched HMM state time series were regressed onto trial-wise decision variables. First, we epoched the HMM state time series either with –1 to +4 s relative to the option onset (“option-locked” analysis) or –1.5 to + 3.9 s relative to the participants’ response (“response-lock” analysis). The epoched time series were baseline corrected using either –0.2 to –0.1 s relative to option onset for the option-locked analysis or +1.5 to +1.6 s relative to feedback onset (so that the baseline fell into the post-trial ITI period) for the response-locked analysis. Next, we assessed the effects of the decision variables on the HMM state time series using the following GLMs:

GLM1 (**Figure 2B**). This model comprised a constant term to estimate the trial-related changes in the HMM state time series, irrespective of the value and choice.

GLM2a (**Figure 3A**). This model comprised two regressors: i) a constant term and ii) the sum of the subjective values of the two options (value sum, *V_chosen_* + *V_unchosen_*). Note that using the value sum as the regressor is mathematically equivalent to using the subjective value of the risky option (*V_risky_*) because the value of the sure option is constant.

GLM2b (**Figure S4A**). This model comprised two regressors: i) a constant term and ii) the reward magnitude of the risky option.

GLM2c (**Figure S4B**). This model comprised two regressors: i) a constant term and ii) the reward probability of the risky option.

GLM3a (**Figure 3B**). This model comprised two regressors: i) a constant term and ii) the difference in subjective values between the two options (value difference, *V_chosen_* – *V_unchosen_*).

GLM3b (**Figure S5AB**). This model comprised two regressors: i) a constant term and ii) the absolute difference in subjective values between the two options (absolute value difference, i.e., |*V_chosen_* – *V_unchosen_*|). Note that the absolute value difference is equivalent to |*V_risky_* – *V_sure_*|, a variable independent of the participant’s choice. We called the sign-flipped version of the variable as value similarity (–|*V_chosen_* – *V_unchosen_*|).

GLM3c (**Figure S5C**). This model comprised two regressors: i) a constant term and ii) trial-wise RT.

GLM4a (**Figure S10A**). This model comprised four regressors: i) a constant term, ii) *V_risky_*, iii) a binary variable denoting the participant’s trial-wise choice (choice type: risky = +1, sure = –1), and iv) an interaction term between *V_risky_* and choice type.

GLM4b (**Figure S10B**). This model comprised two regressors: i) a constant term and ii) *V_risky_*. Notably, we modeled only the trials in which the participants chose risky options. To ensure a reliable estimation at the group level, we included a subset of the participants (*N* = 21) for whom at least 100 trials were available (150.52 ± 23.17 trials, mean ± SD across participants).

GLM4c (**Figure S10C**). This model comprised two regressors: i) a constant term and ii) *V_risky_*. Notably, we modeled only the trials in which the participants chose the sure option. We included all 35 participants because at least 100 trials were available for every participant (170.09 ± 47.68 trials, mean ± SD across participants).

GLM5a (**Figure S11A**). This model comprised four regressors: i) a constant term, ii) *EV_risky_*, iii) choice type, and iv) an interaction term between *EV_risky_* and choice type.

GLM5b (**Figure S11B**). This model comprised four regressors: i) a constant term, ii) *VAR_risky_*, iii) choice type, and iv) an interaction term between *VAR_risky_* and choice type.

GLM6a (**Figure S12A**). This model comprised two regressors: i) a constant term and ii) the subjective value of the outcome (*V_feedback_*) for the risky and sure-choice trials. *V_feedback_* was estimated based on the same utility model (PT) as the other subjective values, where the probability of the revealed outcome was always 100%.

GLM6b (**Figure S12B**). This model comprised two regressors: i) a constant term and ii) reward prediction error (*PE*). *PE* was defined the subjective value of the actual outcome minus the expected subjective value of the possible outcomes.

We estimated a beta coefficient for each participant, state, and time point, resulting in a time series of beta estimates for each participant and state. The corrected beta time series was referred to as task-evoked occupancy, indicating the degree to which the HMM state time series were modulated (across trials) by task variables (Quinn et al., 2018). We performed group-level statistical tests via nonparametric cluster-based permutation implemented in the HMM-MAR toolbox (one-sample *t* test with 1,000 permutations, two tailed, corrected for multiple comparisons across time points and states). Specifically, in each iteration of the permutation, the data from a random subset of the participants were sign-flipped, and the group-level task-evoked occupancy was computed. The null distribution was created by gathering the maximal values of this group-level task-evoked occupancy (across time points and states) over the 1,000 permutations. *P* values were obtained by comparing the actual (observed) maximal values of the group-level task-evoked occupancy with the null distribution.

To investigate frequency-specific whole-brain dynamics, we performed exploratory HMM analyses separately for the 1–4 (delta), 4–8 (theta), 8–13 (alpha), and 15–35 Hz (beta) frequency bands, using the amplitude envelopes in each of these frequency bands instead of the 2–10 Hz frequency band used in the main analysis. All other procedures were identical to those used in the main analysis.

#### Time-resolved MVPA

To examine the robustness of the findings derived from the HMM analysis, we conducted time-resolved MVPA (Liu et al., 2019, Liu et al., 2021, Russek et al., 2024). Both HMM and time-resolved MVPA can be used to probe whole-brain dynamics; however, they differ in several key aspects. In particular, HMM analysis explicitly considers serial dependence across successive time points, whereas time-resolved MVPA treats each time point separately when training and testing a model. Furthermore, HMM analysis does not use trial-wise task variables for state estimation (using them only in the subsequent GLM step), whereas time-resolved MVPA requires trial-wise task variables to tune the model parameters.

For time-resolved MVPA, we used the same parcel-wise 2–10 Hz amplitude envelopes as those used in the HMM analysis. The envelopes were smoothed with a 50-ms moving average filter, z-score normalized for each parcel, epoched from –1 to +4 s relative the option onset for the option-locked analysis and from –1.5 to + 3.9 s relative to the participant’s response for the response-lock analysis, and baseline corrected (–0.2 to –0.1 s relative to option onset for the option-locked analysis and +1.5 to +1.6 s relative to feedback onset for the response-locked analysis). To predict trial-wise decision variables (1 × *N* vector, where *N* is the number of trials) from MEG activity (*M* × *N* feature-by-trial matrix) at each time point in the epoch, we used Least Absolute Shrinkage and Selection Operator (LASSO)-regularized linear regression implemented using *lassoglm* in MATLAB (Liu et al., 2019, Liu et al., 2021, Russek et al., 2024). The model parameters were tuned using a leave-one-run-out nested cross-validation approach. In the outer cross-validation loop, we used three runs as the training set and the left-out run as the testing set. Within each outer-loop training set, we optimized the L1 regularization parameter λ (varying from 0.001 to 0.01 in increments of 0.001) using an inner leave-one-run-out cross-validation loop. We then used the optimal value of λ to train and test the regression model in the outer cross-validation loop. We evaluated prediction performance using Spearman’s correlation coefficient between the actual and predicted values of the trial-wise decision variable (across all *N* trials). Correlation coefficients were Fisher Z-transformed before group-level analysis. By repeating this procedure over time points, we obtained a time series of the prediction performance. We performed group-level statistical tests using nonparametric cluster-based permutation in the same manner as in the HMM analysis (one-sample t test with 1,000 permutations, corrected for time points). To obtain a two-dimensional temporal generalization matrix, we repeated the same procedure across all pairs of training and testing time points, using three runs for training and the left-out run for testing.

#### Source-level GLM analysis

In addition to the HMM analysis, we performed a conventional source-level GLM analysis to localize simple visual- and motor-related brain responses. We used the same source-reconstructed MEG signals (2–10 Hz) as in the HMM analysis, but retained the grid-wise time series. We used the same definitions of epochs and baseline periods as in the HMM analysis and ran a GLM that modeled a constant term for each participant, grid, and time point. Group-level *t* values (one-sample *t* tests) were computed for each grid and time point.

#### Time-frequency analysis

We also performed a time-frequency analysis in brain regions that were previously implicated in value-based decision making. First, we defined a set of ROIs based on previous literature (Levy and Glimcher, 2012, Bartra et al., 2013, Clithero and Rangel, 2014, Levy, 2017): the anterior striatum, anterior insula, preSMA, precuneus, vmPFC, and posterior parietal cortex (PPC). We used the Automated Anatomical Labeling (Tzourio-Mazoyer et al., 2002) to define the anterior striatum (bilateral caudate and putamen, y ≥ 0) (Izuma et al., 2010) and anterior insula (bilateral insula, y ≥ 0) (Hölzel et al., 2008, Morelli and Lieberman, 2013). For the preSMA, precuneus, ventromedial prefrontal cortex, and parietal lobule, we defined the ROIs as the clusters reported in the meta-analysis (Corlett et al., 2022), because the AAL was not suitable for defining these regions (too large or ill-split by anatomical borders). Although the ROIs derived from Corlett et al. (2022) pertain to prediction error and not specifically to subjective value, these regions empirically matched with regions reported in studies of value-based decision making (Wunderlich et al., 2009, Hare et al., 2011, Levy and Glimcher, 2012, Clithero and Rangel, 2014). Next, we extracted the ROI-wise time series from the source-reconstructed MEG time series as the first principal component derived from PCA. We performed a wavelet analysis (with a morlet factor of 6) between 1 and 45 Hz with 2-Hz frequency bins. We set the time window of interest as −0.6 to +3.6 s with respect to the option onset in the option-locked analysis and −1.1 to +3.5 s with respect to the participants’ response in the response-locked analysis, which excluded the initial and last 0.4-s periods from the original epochs used in the HMM and source-level GLM analysis to avoid the edge effect. We then performed GLM analyses in each ROI and frequency bin, in the same manner as we did in the HMM analysis.

## Acknowledgements

We thank K. Matsumori, H. Takeichi, N. Hironaga, K. Amano, A. Gunji, T. Sumiyoshi, H. Takahashi, Y. Kaneko, and K. Nakagome for helpful discussions. This study was supported by JST [Moonshot R&D][Grant Number JPMJMS2294], MEXT Grant-in-Aid for Scientific Research on Innovative Areas [Grant Number JP21H00222] awarded to M. Matsumoto, and JSPS [KAKENHI] [Grant Number JP17H05929] awarded to K. Iijima.

## Figures

**Figure S1.**
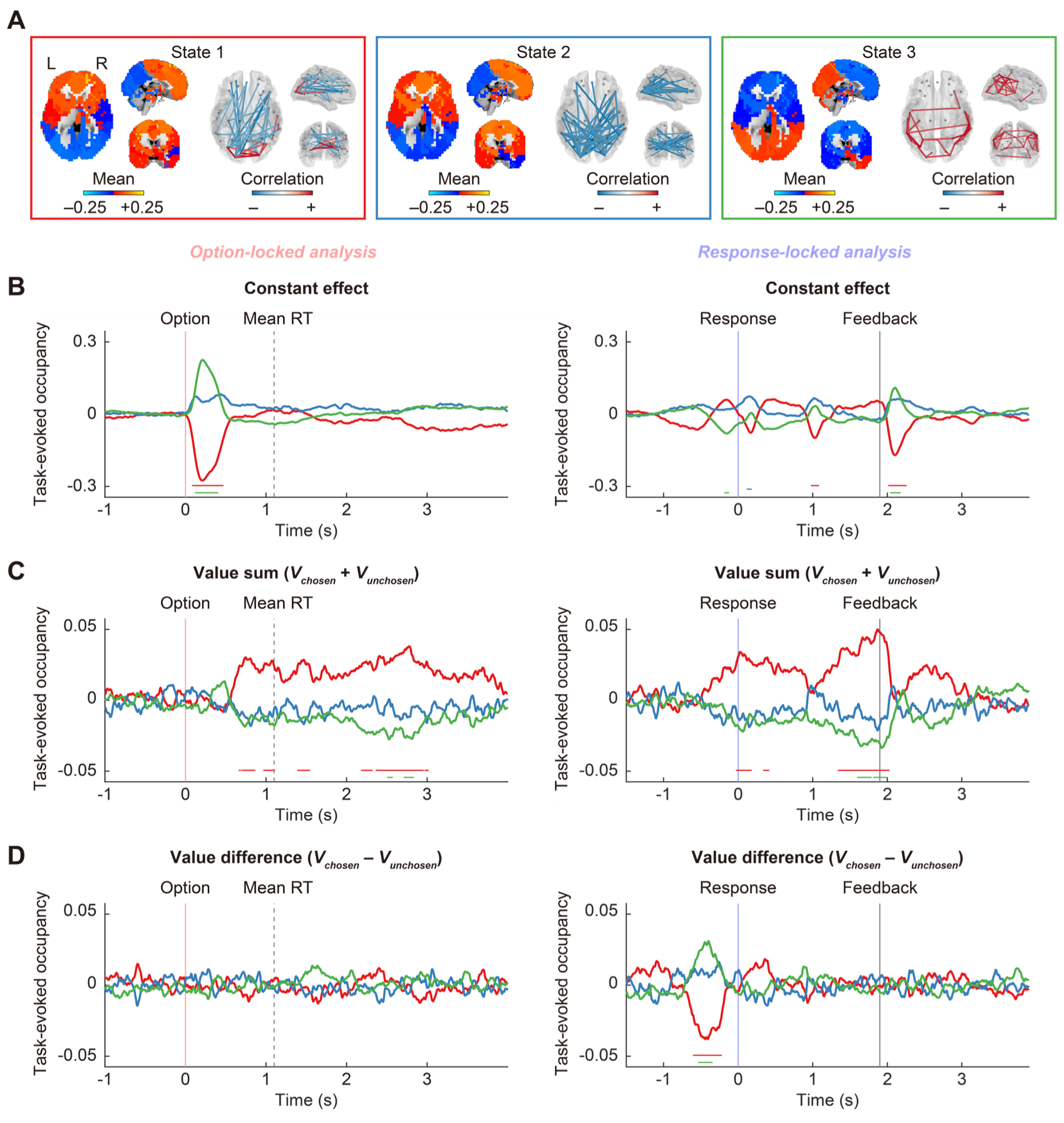
Task-evoked occupancy derived from HMM analysis with number of states (*K*) = 3. **A** Whole-brain pattern of each HMM state characterized by parcel-wise means and inter-parcel correlations of amplitude envelopes. The mean values are derived from parcel-wise z-scored amplitude envelopes averaged across time points within each state. Correlations (edges) are depicted for parcel pairs exhibiting top 5% strength in absolute value (scaled by maximum absolute value for illustration, which is invariant with respect to 0). R, right hemisphere; L, left hemisphere. **B** Task-evoked occupancy associated with the constant effect across trials. **C** Task-evoked occupancy associated with value sum (*V_chosen_* + *V_unchosen_*). **D** Task-evoked occupancy associated with value difference (*V_chosen_* – *V_unchosen_*). Each time series corresponds to each state shown in Panel A with the same color (red: state 1, blue: state 2, green: state 3). Horizontal lines at the bottom indicate *P* < 0.05, cluster-based permutation test. The left and right panels show the results of option-locked and response-locked analyses, respectively.

**Figure S2.**
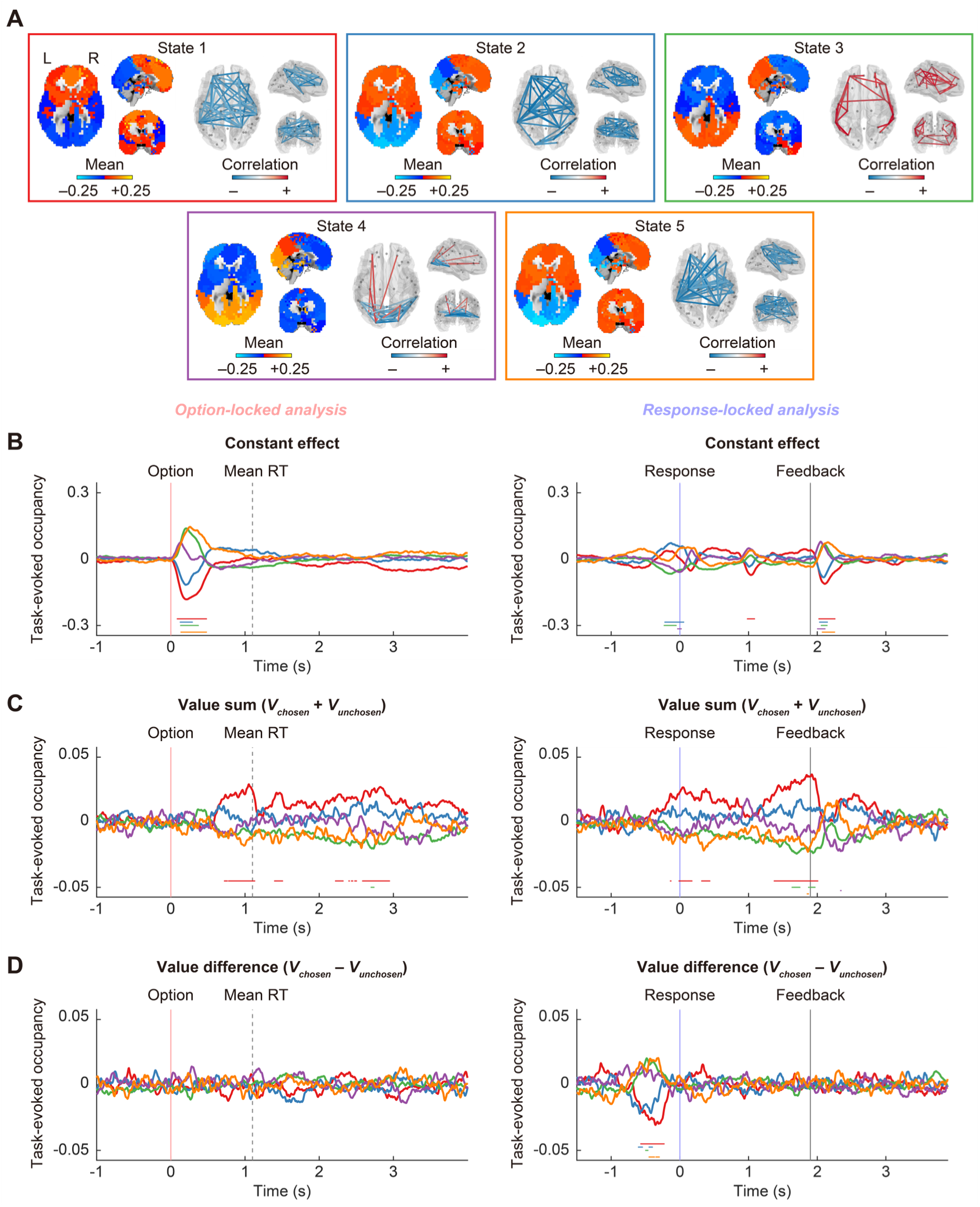
Task-evoked occupancy derived from HMM analysis with number of states (*K*) = 5. **A** Whole-brain pattern of each HMM state characterized by parcel-wise means and inter-parcel correlations of amplitude envelopes. The mean values are derived from parcel-wise z-scored amplitude envelopes averaged across time points within each state. Correlations (edges) are depicted for parcel pairs exhibiting top 5% strength in absolute value (scaled by maximum absolute value for illustration, which is invariant with respect to 0). R, right hemisphere; L, left hemisphere. **B** Task-evoked occupancy associated with the constant effect across trials. **C** Task-evoked occupancy associated with value sum (*V_chosen_* + *V_unchosen_*). **D** Task-evoked occupancy associated with value difference (*V_chosen_* – *V_unchosen_*). Each time series corresponds to each state shown in Panel A with the same color (red: state 1, blue: state 2, green: state 3, purple: state 4, orange: state 5). Horizontal lines at the bottom indicate *P* < 0.05, cluster-based permutation test. The left and right panels show the results of option-locked and response-locked analyses, respectively.

**Figure S3.**
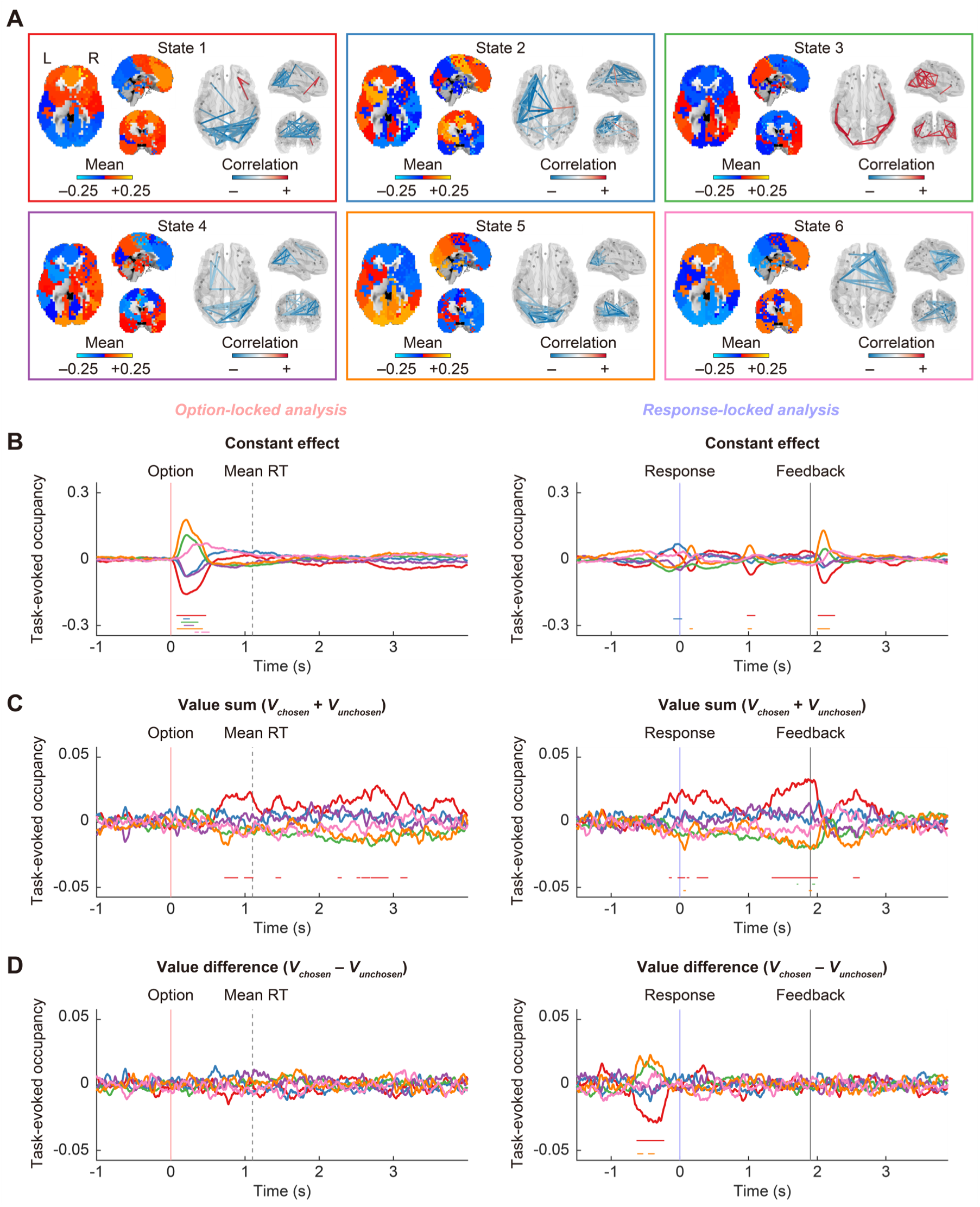
Task-evoked occupancy derived from HMM analysis with number of states (*K*) = 6. **A** Whole-brain pattern of each HMM state characterized by parcel-wise means and inter-parcel correlations of amplitude envelopes. The mean values are derived from parcel-wise z-scored amplitude envelopes averaged across time points within each state. Correlations (edges) are depicted for parcel pairs exhibiting top 5% strength in absolute value (scaled by maximum absolute value for illustration, which is invariant with respect to 0). R, right hemisphere; L, left hemisphere. **B** Task-evoked occupancy associated with the constant effect across trials. **C** Task-evoked occupancy associated with value sum (*V_chosen_* + *V_unchosen_*). **D** Task-evoked occupancy associated with value difference (*V_chosen_* – *V_unchosen_*). Each time series corresponds to each state shown in Panel A with the same color (red: state 1, blue: state 2, green: state 3, purple: state 4, orange: state 5, pink: state 6). Horizontal lines at the bottom indicate *P* < 0.05, cluster-based permutation test. The left and right panels show the results of option-locked and response-locked analyses, respectively.

**Figure S4.**
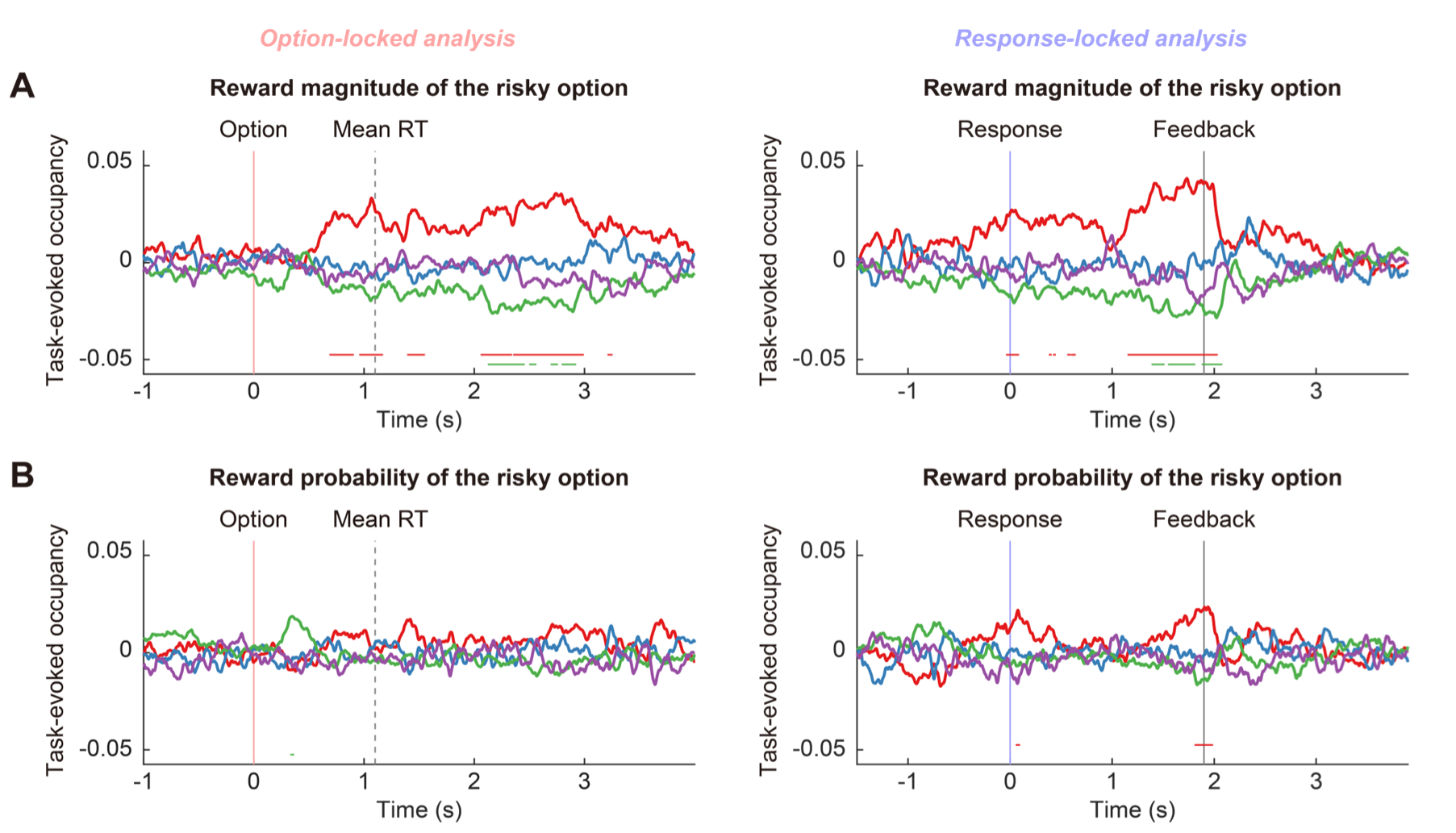
Task-evoked occupancy associated with reward magnitude and probability. **A** Task-evoked occupancy associated with reward magnitude of the risky option (ranging from 300 to 5000 yen). **B** Task-evoked occupancy associated with reward probability of the risky option (ranging from 13 to 75%). The colors of the time series correspond to the HMM states presented in Figure 2A (red: state 1, blue: state 2, green: state 3, purple: state 4). Horizontal lines at the bottom indicate *P* < 0.05, cluster-based permutation test. The left and right panels show the results of option-locked and response-locked analyses, respectively.

**Figure S5.**
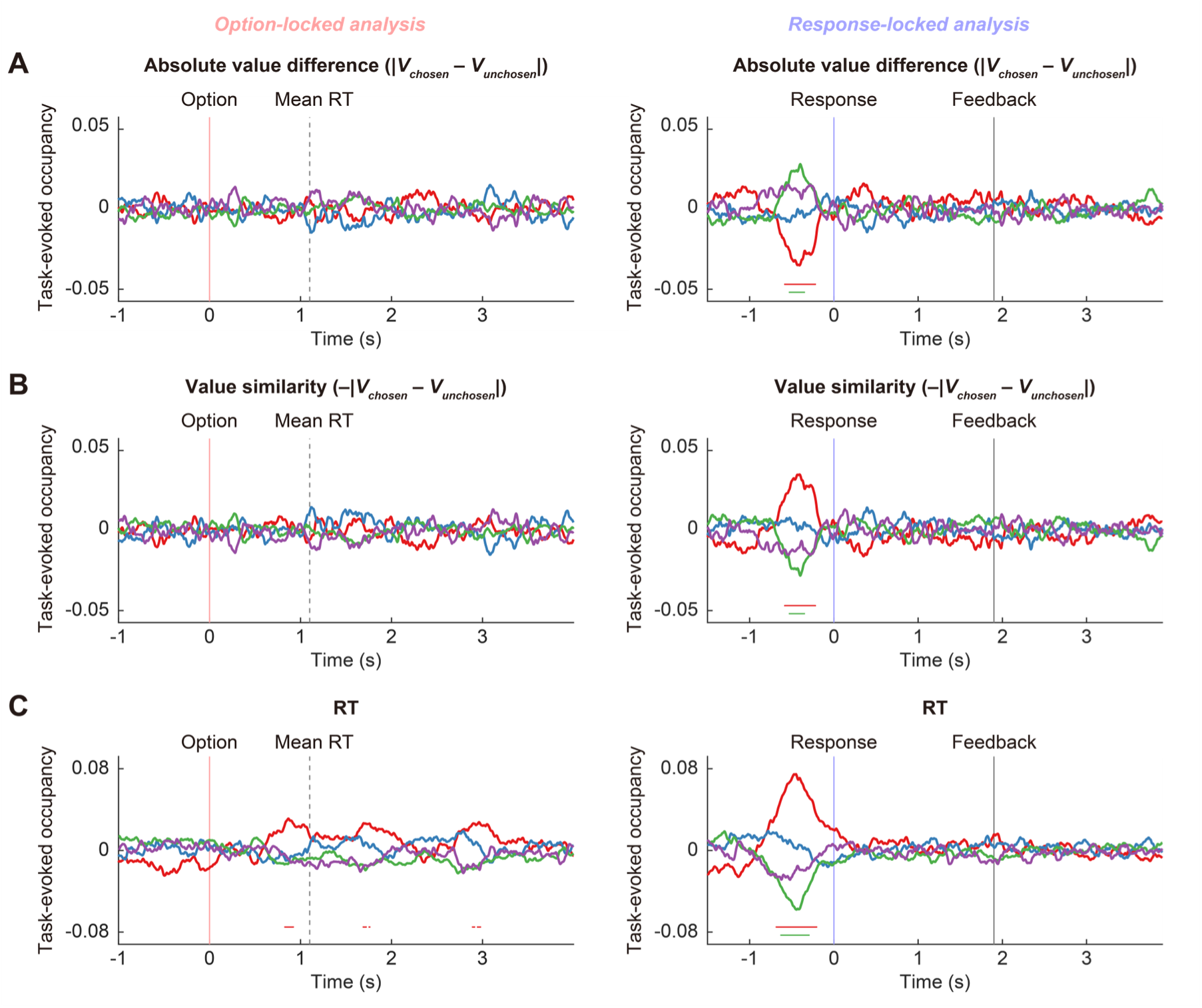
Task-evoked occupancy associated with variables reflecting value difference. **A** Task-evoked occupancy associated with absolute value difference (|*V_chosen_* – *V_unchosen_*|). **B** Task-evoked occupancy associated with value similarity (–|*V_chosen_* – *V_unchosen_*|). Note that value similarity is defined as the absolute value difference multiplied by –1, therefore panels A and B show the same results (except that time series in panel B are flipped with respect to *y* = 0 compared to those in panel A to facilitate comparison with results associated with RT). **C** Task-evoked occupancy associated with RT. The colors of the time series correspond to the HMM states presented in Figure 2A (red: state 1, blue: state 2, green: state 3, purple: state 4). Horizontal lines at the bottom indicate *P* < 0.05, cluster-based permutation test. The left and right panels show the results of option-locked and response-locked analyses, respectively.

**Figure S6.**
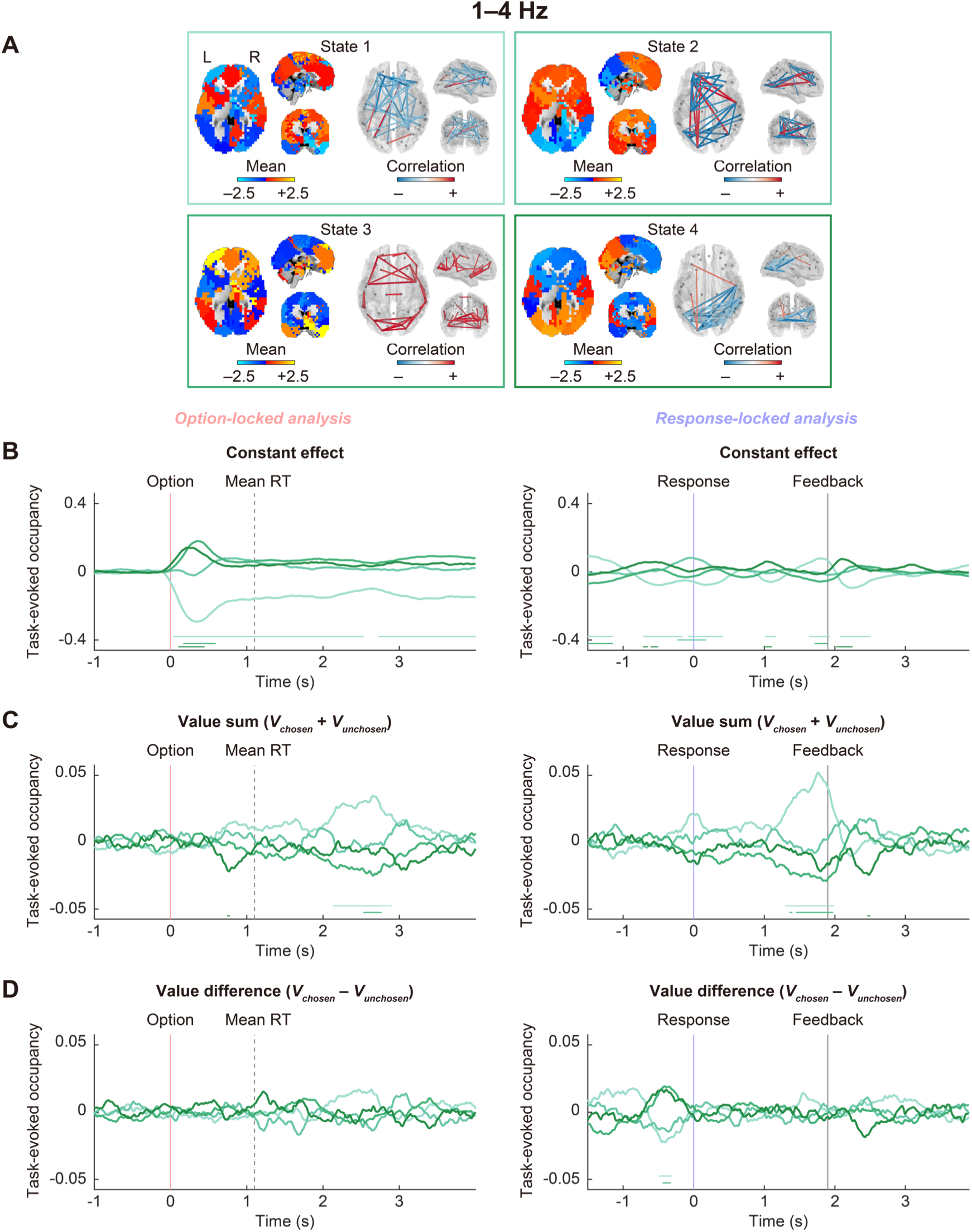
Task-evoked occupancy derived from HMM analysis in 1–4 Hz frequency band. **A** Whole-brain pattern of each HMM state characterized by parcel-wise means and inter-parcel correlations of amplitude envelopes. The mean values are derived from parcel-wise z-scored amplitude envelopes averaged across time points within each state. Correlations (edges) are depicted for parcel pairs exhibiting top 5% strength in absolute value (scaled by maximum absolute value for illustration, which is invariant with respect to 0). R, right hemisphere; L, left hemisphere. **B** Task-evoked occupancy associated with the constant effect across trials. **C** Task-evoked occupancy associated with value sum (*V_chosen_* + *V_unchosen_*). **D** Task-evoked occupancy associated with value difference (*V_chosen_* – *V_unchosen_*). Each time series corresponds to each state shown in Panel A with the same color. Horizontal lines at the bottom indicate P < 0.05, cluster-based permutation test. The left and right panels show the results of option-locked and response-locked analyses, respectively.

**Figure S7.**
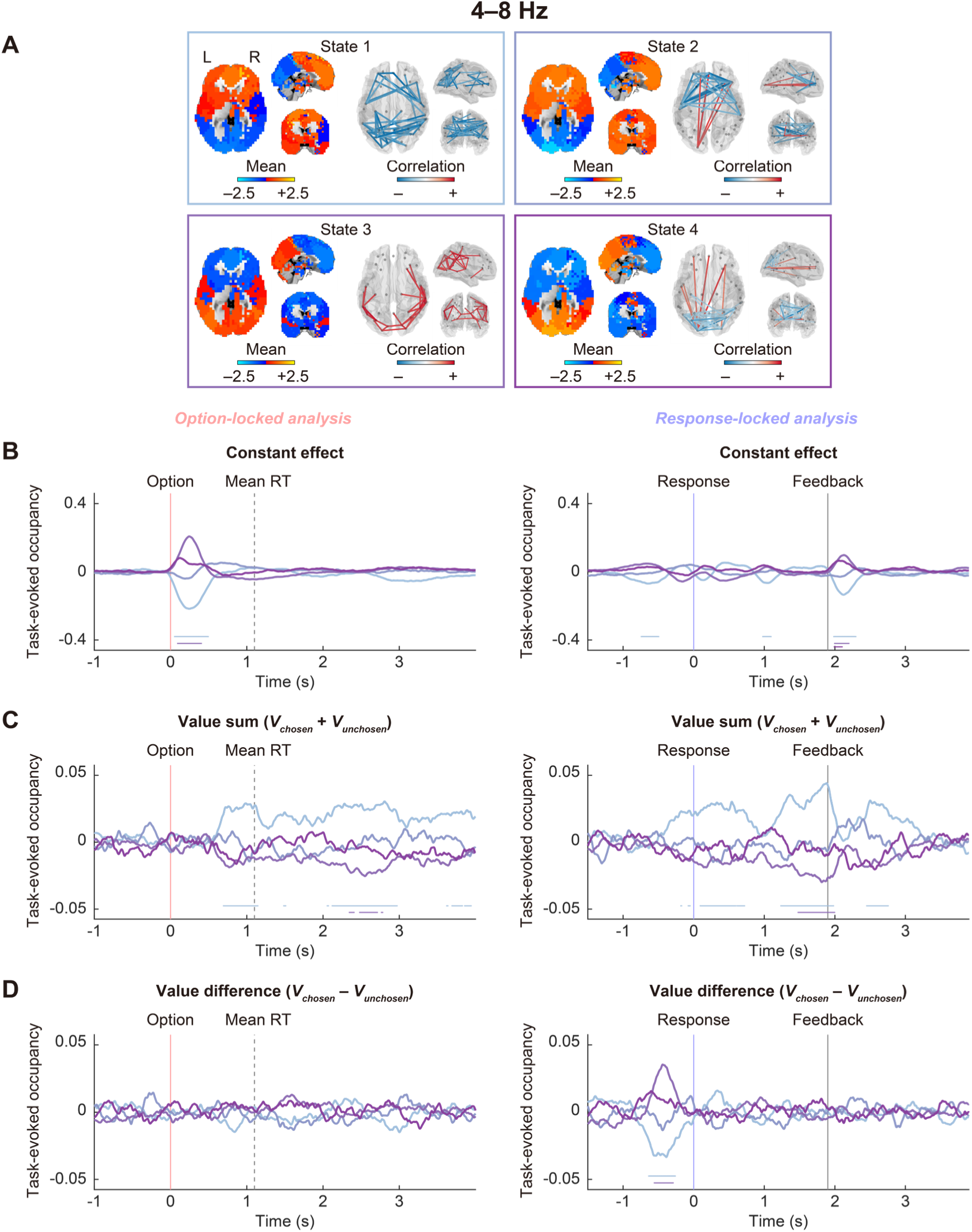
Task-evoked occupancy derived from HMM analysis in 4–8 Hz frequency band. **A** Whole-brain pattern of each HMM state characterized by parcel-wise means and inter-parcel correlations of amplitude envelopes. The mean values are derived from parcel-wise z-scored amplitude envelopes averaged across time points within each state. Correlations (edges) are depicted for parcel pairs exhibiting top 5% strength in absolute value (scaled by maximum absolute value for illustration, which is invariant with respect to 0). R, right hemisphere; L, left hemisphere. **B** Task-evoked occupancy associated with the constant effect across trials. **C** Task-evoked occupancy associated with value sum (*V_chosen_* + *V_unchosen_*). **D** Task-evoked occupancy associated with value difference (*V_chosen_* – *V_unchosen_*). Each time series corresponds to each state shown in Panel A with the same color. Horizontal lines at the bottom indicate P < 0.05, cluster-based permutation test. The left and right panels show the results of option-locked and response-locked analyses, respectively.

**Figure S8.**
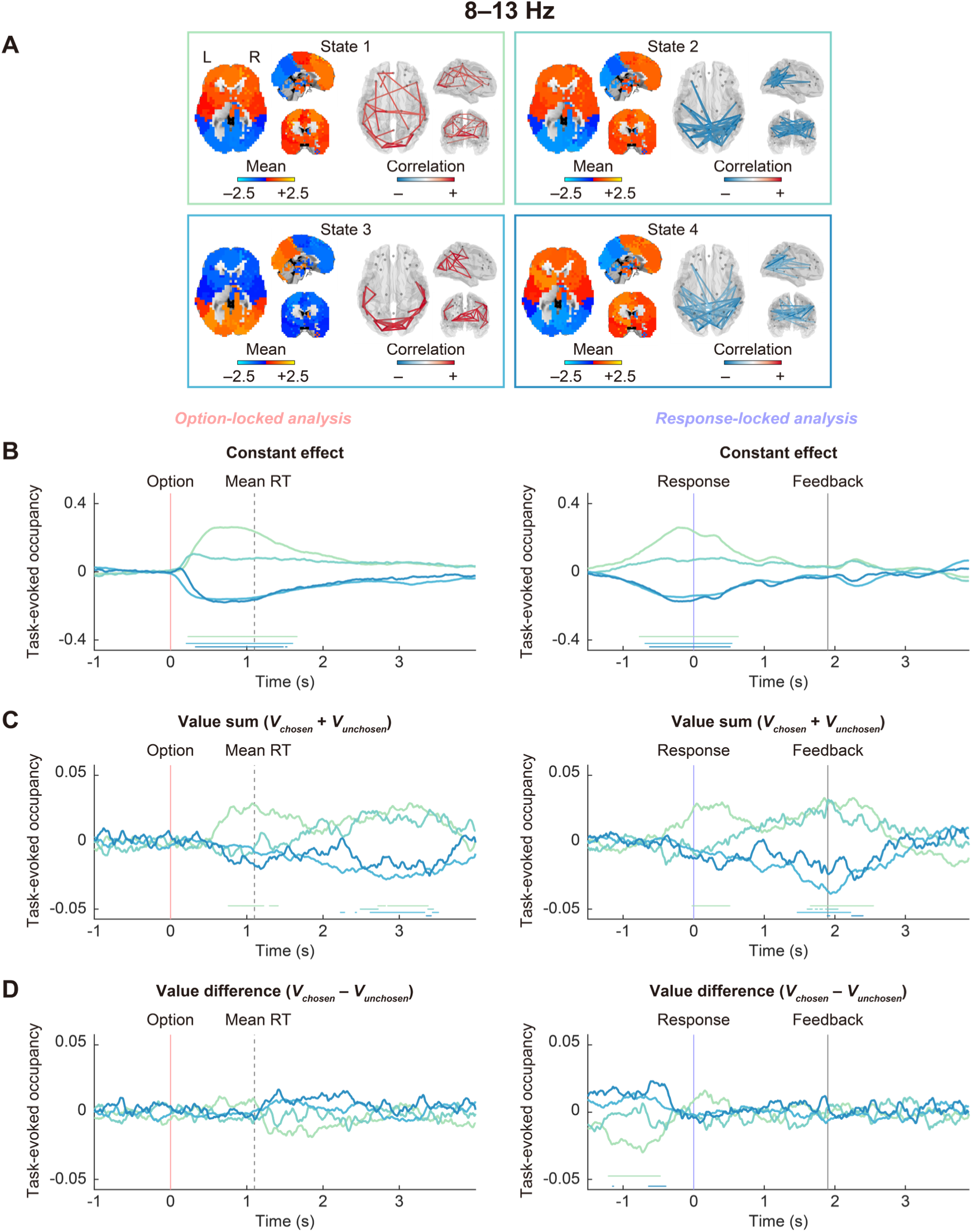
Task-evoked occupancy derived from HMM analysis in 8–13 Hz frequency band. **A** Whole-brain pattern of each HMM state characterized by parcel-wise means and inter-parcel correlations of amplitude envelopes. The mean values are derived from parcel-wise z-scored amplitude envelopes averaged across time points within each state. Correlations (edges) are depicted for parcel pairs exhibiting top 5% strength in absolute value (scaled by maximum absolute value for illustration, which is invariant with respect to 0). R, right hemisphere; L, left hemisphere. **B** Task-evoked occupancy associated with the constant effect across trials. **C** Task-evoked occupancy associated with value sum (*V_chosen_* + *V_unchosen_*). **D** Task-evoked occupancy associated with value difference (*V_chosen_* – *V_unchosen_*). Each time series corresponds to each state shown in Panel A with the same color. Horizontal lines at the bottom indicate P < 0.05, cluster-based permutation test. The left and right panels show the results of option-locked and response-locked analyses, respectively.

**Figure S9.**
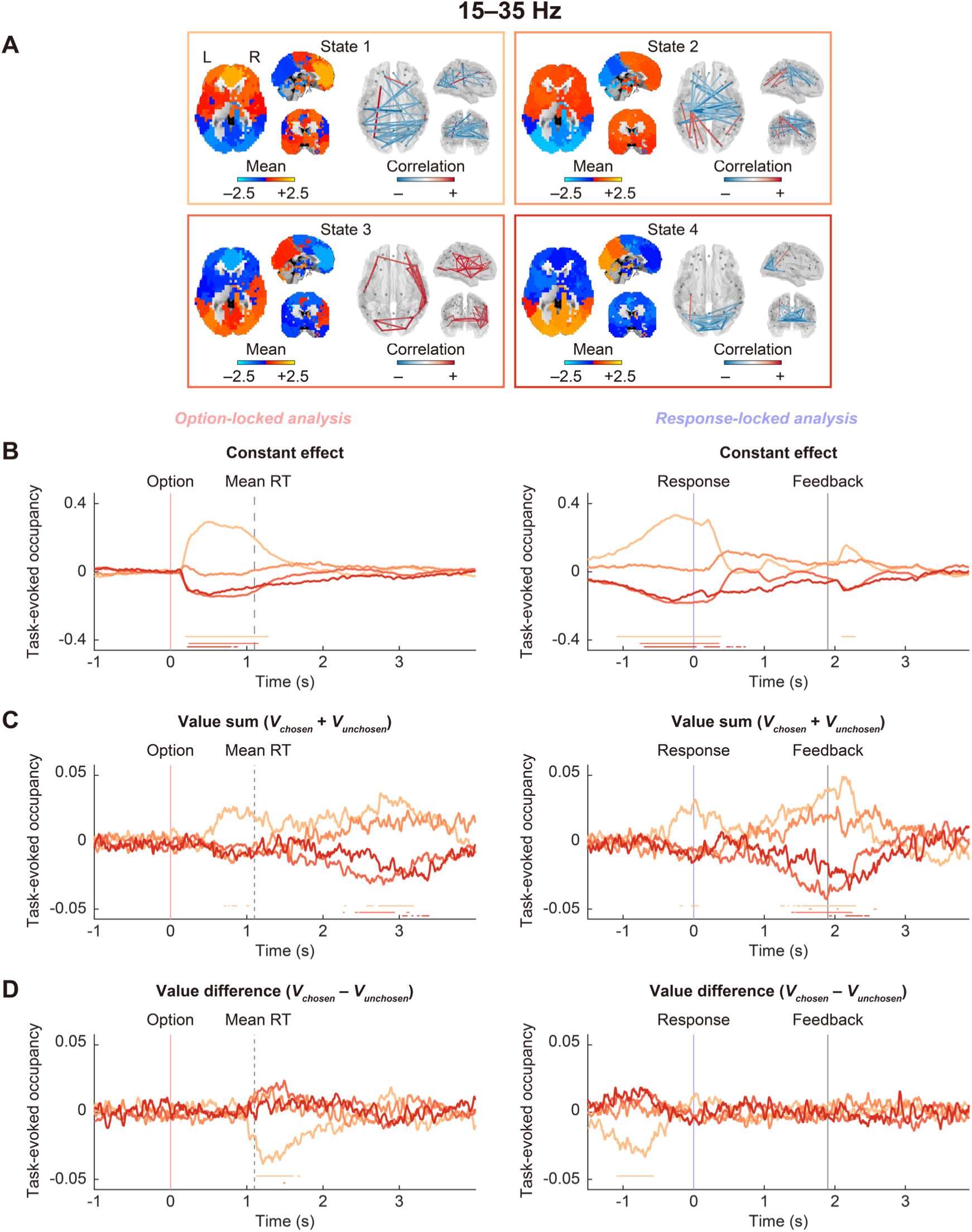
Task-evoked occupancy derived from HMM analysis in 15–35 Hz frequency band. **A** Whole-brain pattern of each HMM state characterized by parcel-wise means and inter-parcel correlations of amplitude envelopes. The mean values are derived from parcel-wise z-scored amplitude envelopes averaged across time points within each state. Correlations (edges) are depicted for parcel pairs exhibiting top 5% strength in absolute value (scaled by maximum absolute value for illustration, which is invariant with respect to 0). R, right hemisphere; L, left hemisphere. **B** Task-evoked occupancy associated with the constant effect across trials. **C** Task-evoked occupancy associated with value sum (*V_chosen_* + *V_unchosen_*). **D** Task-evoked occupancy associated with value difference (*V_chosen_* – *V_unchosen_*). Each time series corresponds to each state shown in Panel A with the same color. Horizontal lines at the bottom indicate P < 0.05, cluster-based permutation test. The left and right panels show the results of option-locked and response-locked analyses, respectively.

**Figure S10.**
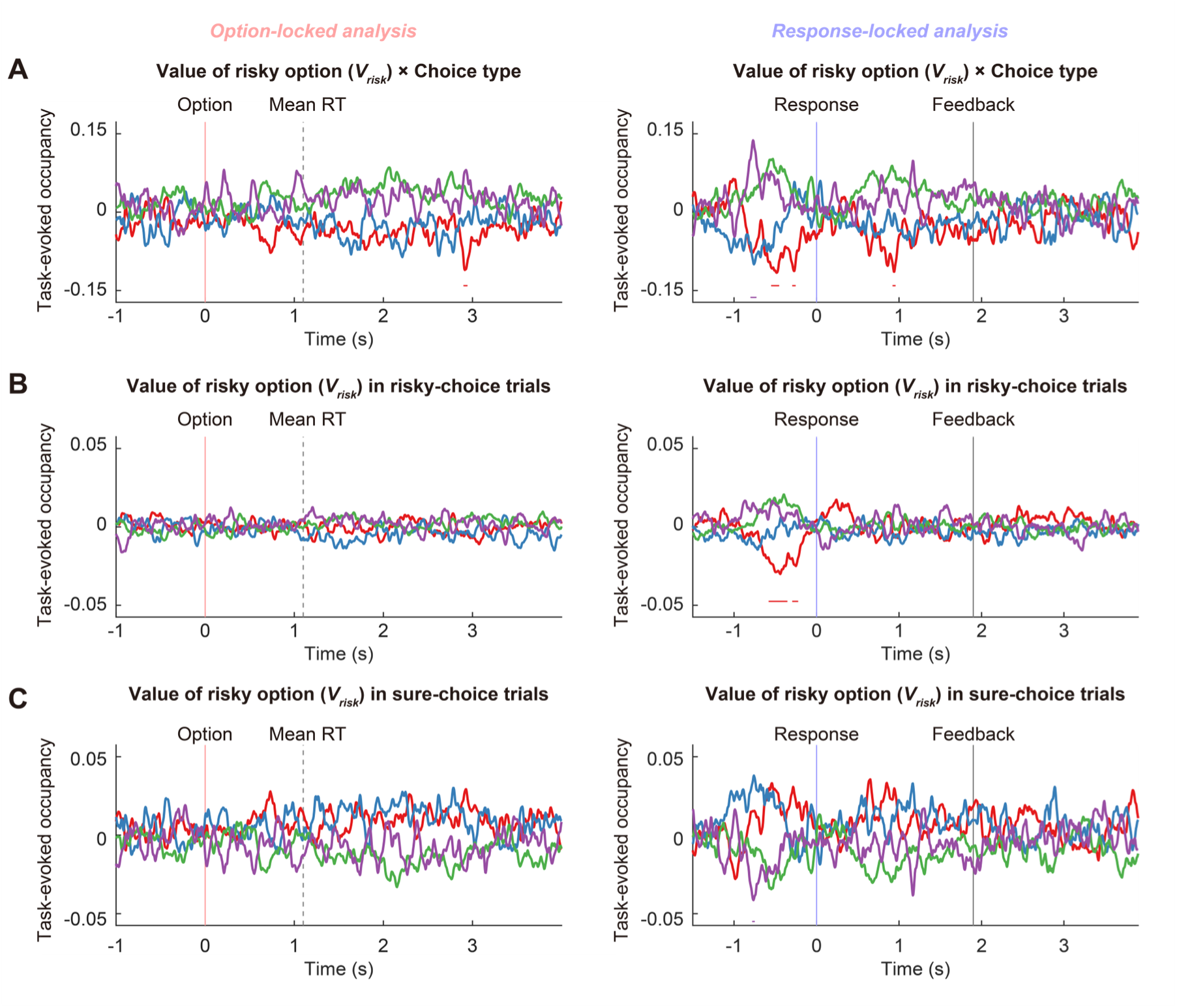
Task-evoked occupancy differentiated by participant’s choice. **A** Task-evoked occupancy associated with the interaction between value of risky option (*V_risky_*) and choice type (risky choice: +1, sure choice: –1). **B** Task-evoked occupancy associated with *V_risky_* in risky-choice trials. **C** Task-evoked occupancy associated with *V_risky_* in sure-choice trials. The colors of the time series correspond to the HMM states presented in Figure 2A (red: state 1, blue: state 2, green: state 3, purple: state 4). Horizontal lines at the bottom indicate *P* < 0.05, cluster-based permutation test. The left and right panels show the results of option-locked and response-locked analyses, respectively.

**Figure S11.**
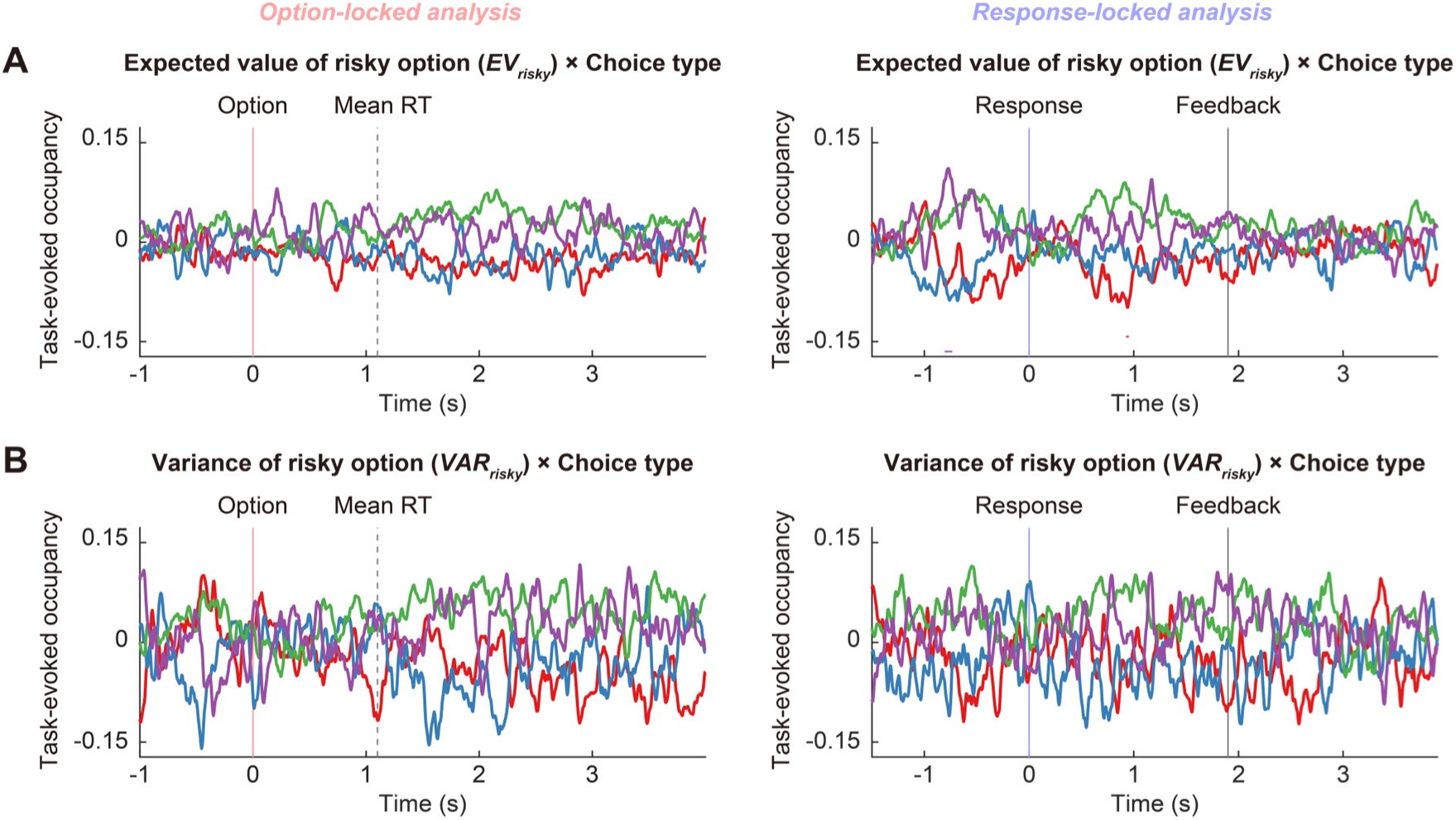
Task-evoked occupancy associated with expected value and variance of risky option. **A** Task-evoked occupancy associated with the interaction between expected value of risky option (*EV_risky_*, i.e., *r* × *p*) and choice type (risky choice: +1, sure choice: –1). **B** Task-evoked occupancy associated with the interaction between variance of risky option (*VAR_risky_*, i.e., *r*^2^ × *p* × (1 – *p*)) and choice type. The colors of the time series correspond to the HMM states presented in Figure 2A (red: state 1, blue: state 2, green: state 3, purple: state 4). Horizontal lines at the bottom indicate *P* < 0.05, cluster-based permutation test. The left and right panels show the results of option-locked and response-locked analyses, respectively.

**Figure S12.**
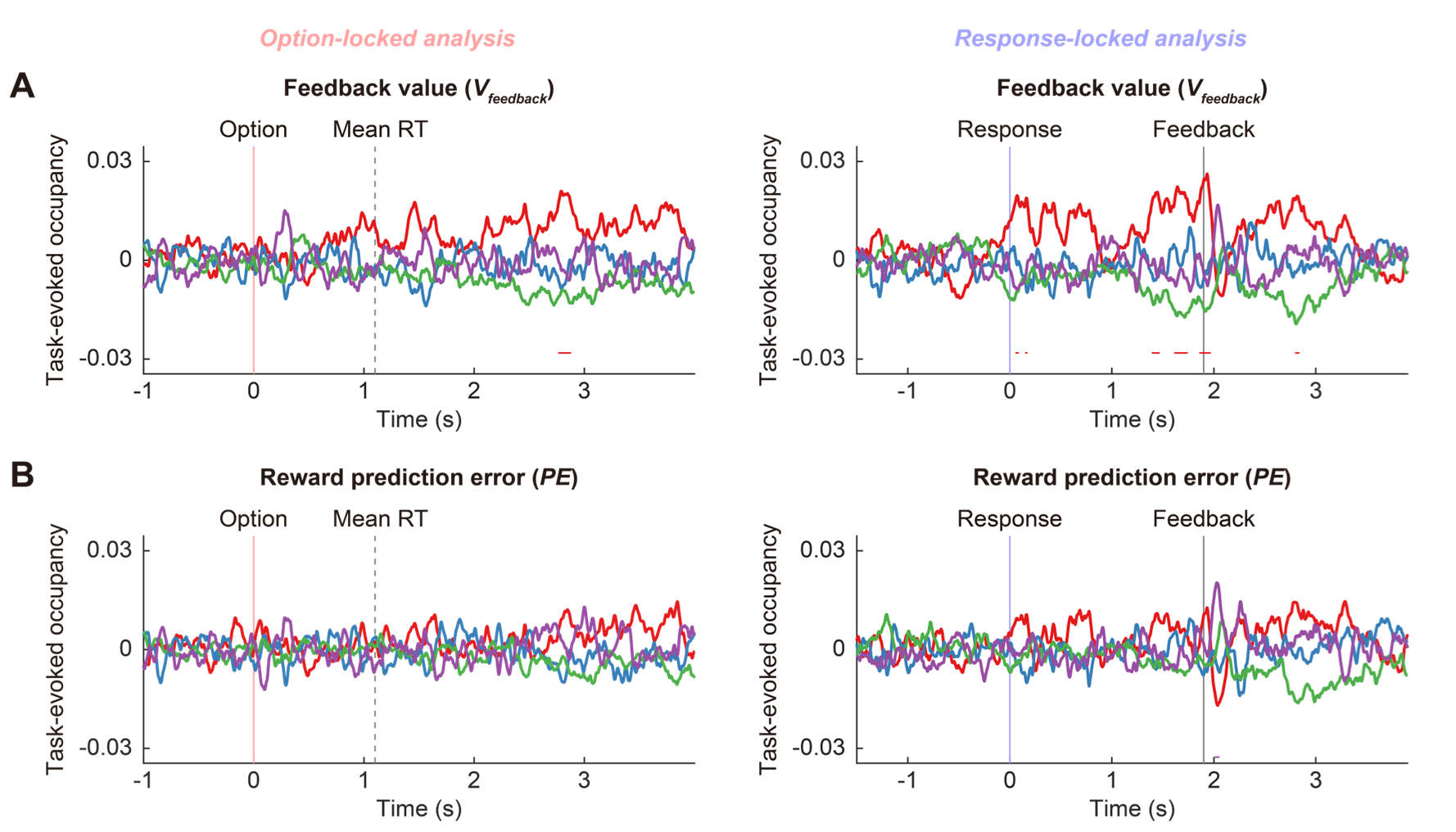
Task-evoked occupancy associated with outcome feedback. **A** Task-evoked occupancy associated with subjective value of outcome feedback (*V_feedback_*). **B** Task-evoked occupancy associated with reward prediction error (*PE*). The colors of the time series correspond to the HMM states presented in Figure 2A (red: state 1, blue: state 2, green: state 3, purple: state 4). Horizontal lines at the bottom indicate *P* < 0.05, cluster-based permutation test. The left and right panels show the results of option-locked and response-locked analyses, respectively.

**Figure S13.**
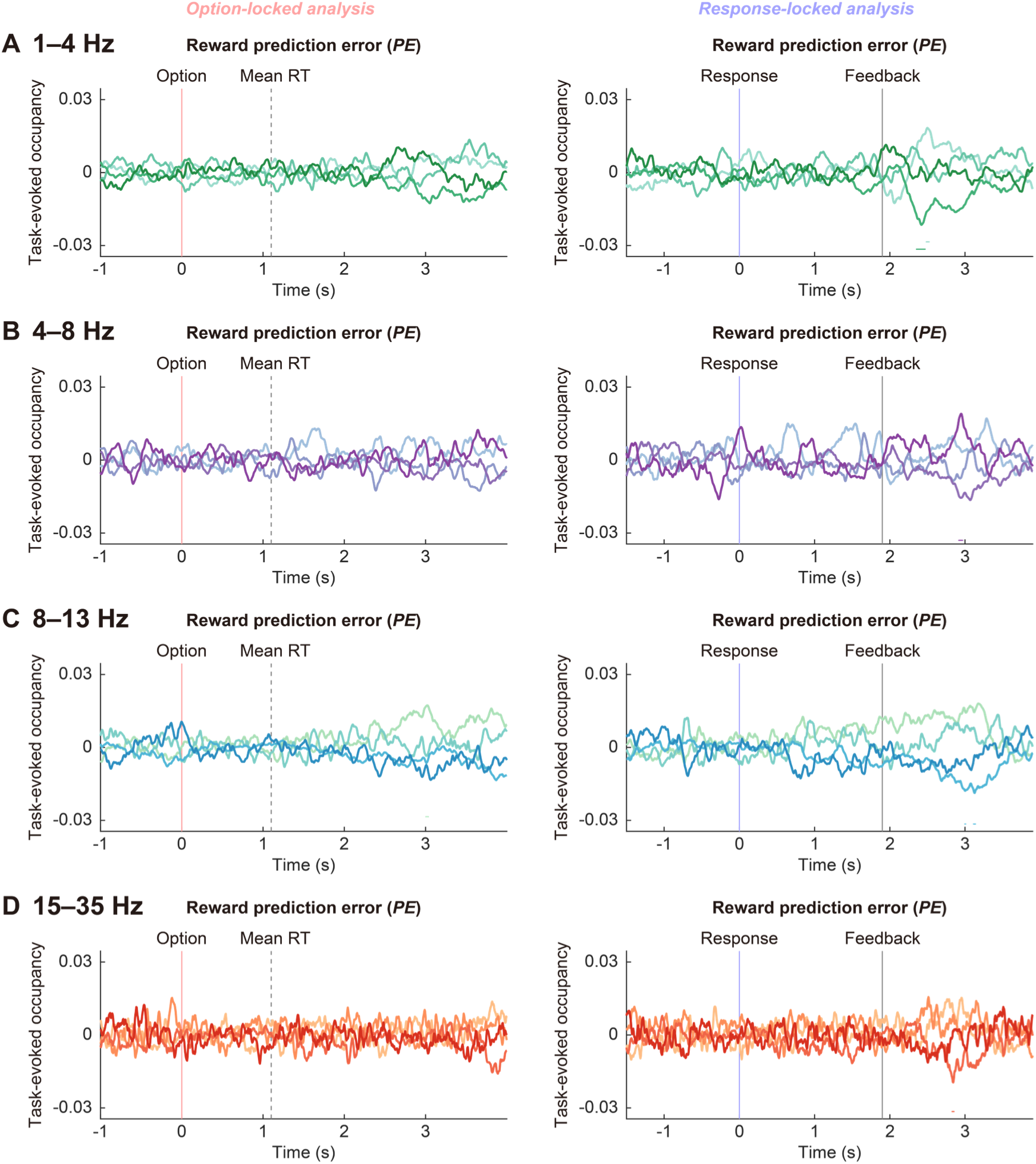
Task-evoked occupancy associated with reward prediction error in different frequency bands. **A** Task-evoked occupancy associated with reward prediction error (*PE*) in the 1–4 Hz frequency band. **B** Task-evoked occupancy associated with *PE* in the 4–8 Hz frequency band. **C** Task-evoked occupancy associated with *PE* in the 8–13 Hz frequency band. **D** Task-evoked occupancy associated with *PE* in the 15–35 Hz frequency band. The colors of the time series correspond to the HMM states presented in Figures S6–S9. Horizontal lines at the bottom indicate *P* < 0.05, cluster-based permutation test. The left and right panels show the results of option-locked and response-locked analyses, respectively.

**Figure S14.**
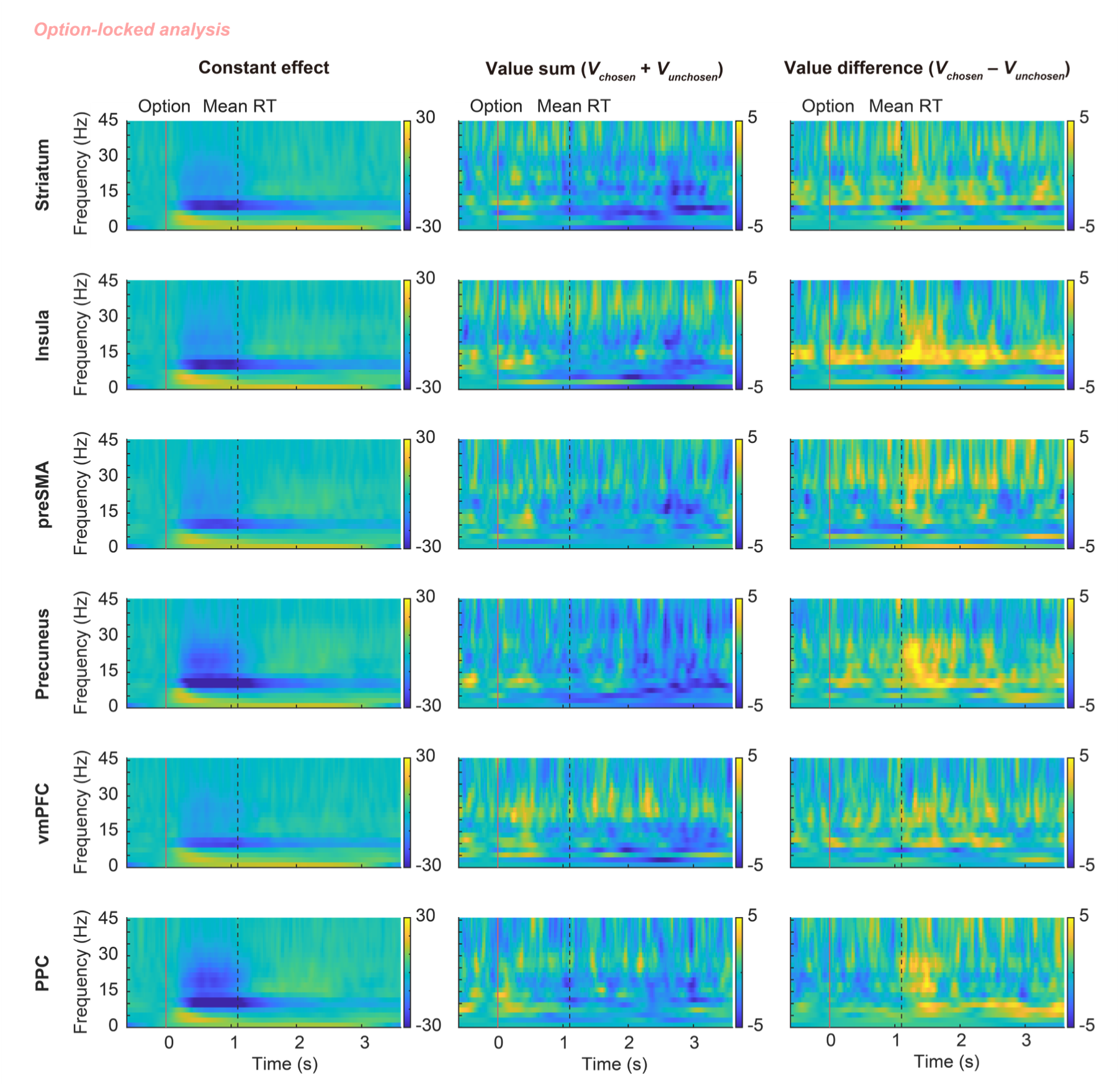
Time frequency representations derived from option-locked analysis. ROI-wise time-frequency representations showing the constant effect across trials (left column), value sum (middle column), and value difference (right column). preSMA, pre-supplementary motor area; vmPFC, ventromedial prefrontal cortex; PPC, posterior parietal cortex.

**Figure S15.**
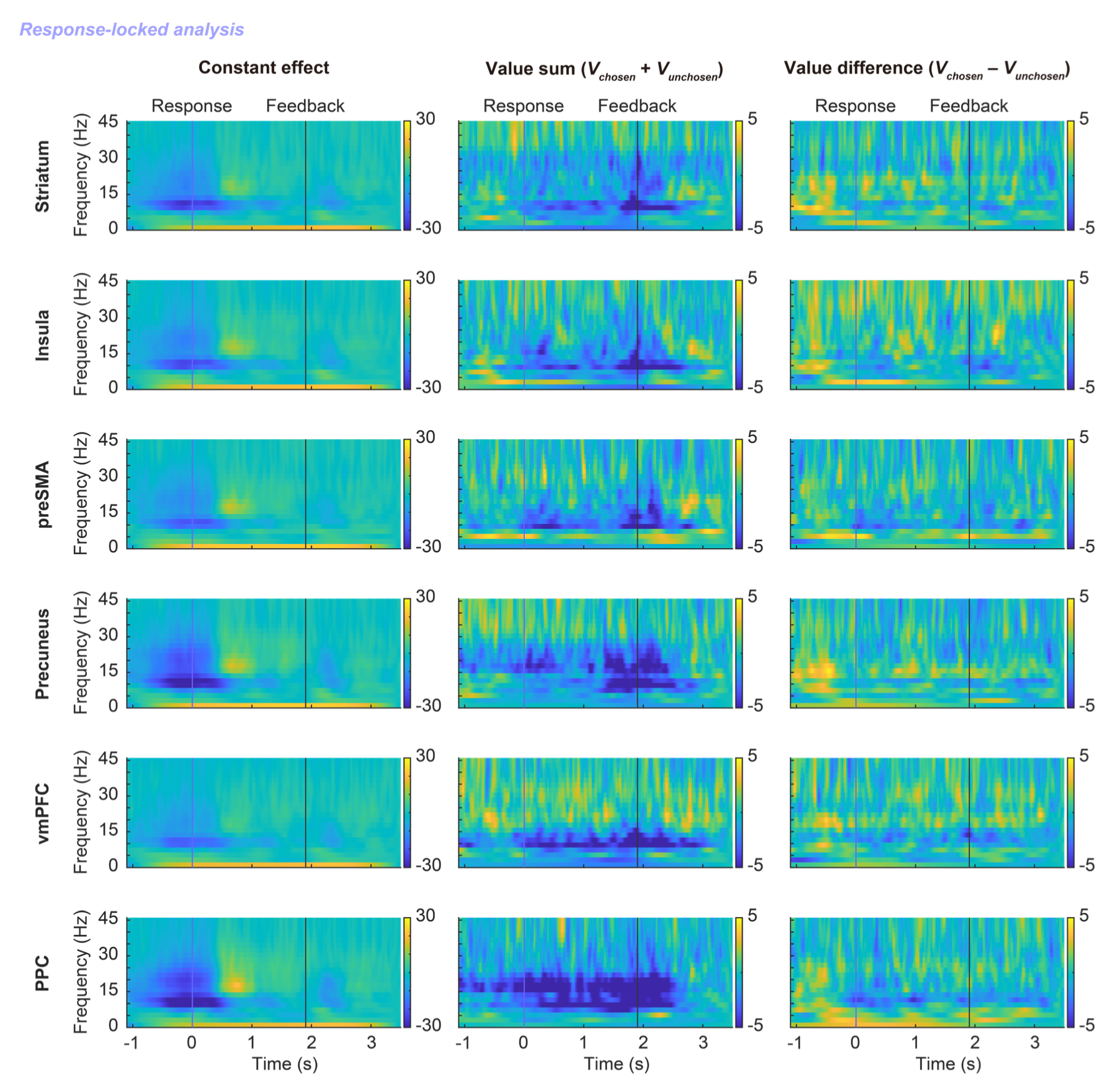
Time frequency representations derived from response-locked analysis. ROI-wise time-frequency representations showing the constant effect across trials (left column), value sum (middle column), and value difference (right column). preSMA, pre-supplementary motor area; vmPFC, ventromedial prefrontal cortex; PPC, posterior parietal cortex.

**Figure S16.**
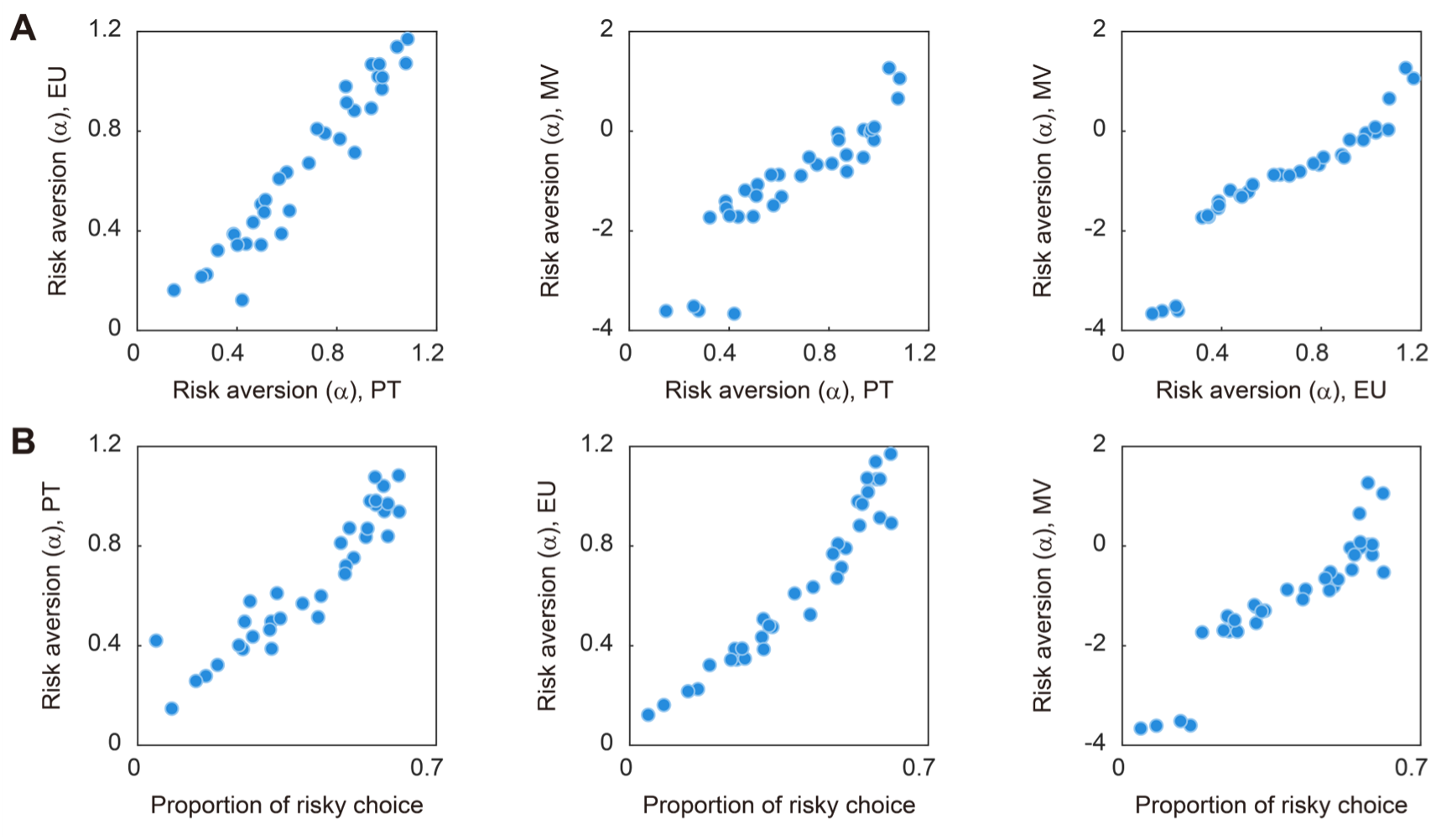
Risk aversion parameters estimated by different behavioral models. **A** Across-participant correlations among risk aversion parameters estimated by prospect theory (RT), exponential utility (EU), and mean-variance (MV) models. **B** Across-participant correlations between proportion of risky choice and risk aversion parameters derived from PT, EU, and MV models.

